# Spatial enhancer activation determines inhibitory neuron identity

**DOI:** 10.1101/2023.01.30.525356

**Authors:** Elena Dvoretskova, May C. Ho, Volker Kittke, Florian Neuhaus, Ilaria Vitali, Daniel D. Lam, Irene Delgado, Chao Feng, Miguel Torres, Juliane Winkelmann, Christian Mayer

## Abstract

The mammalian telencephalon contains a tremendous diversity of GABAergic projection neuron and interneuron types, that originate in a germinal zone of the embryonic basal ganglia. How genetic information in this transient structure is transformed into different cell types is not yet fully understood. Using a combination of *in vivo* CRISPR perturbation, lineage tracing, and ChIP-seq in mice, we found that the transcription factor MEIS2 favors the development of projection neurons through genomic binding sites in regulatory enhancers of projection neuron specific genes. MEIS2 requires the presence of the homeodomain transcription factor DLX5 to direct its functional activity towards these sites. In interneuron precursors, the activation of projection neuron specific enhancers by MEIS2 and DLX5 is repressed by the transcription factor LHX6. When MEIS2 carries a mutation associated with intellectual disability in humans, it is less effective at activating enhancers involved in projection neuron development. This suggests that GABAergic differentiation may be impaired in patients carrying this mutation. Our research supports a model (“Differential Binding‘) where the spatial specific composition of transcription factors at *cis*-regulatory elements determines differential gene expression and cell fate decisions in the ganglionic eminence.

## Introduction

The ganglionic eminences (GEs) are embryonic subpallidal structures that give rise to numerous GABAergic inhibitory cell types (Bandler et al., 2017). It is divided into three spatial regions: the medial (MGE), caudal (CGE), and lateral (LGE) GEs (Wonders and Anderson, 2006; Gelman et al., 2011; Anderson et al., 2001). For example, the MGE and CGE produce many distinct types of interneurons of the cortex, striatum, and hippocampus (Butt et al., 2005; Nery et al., 2002; Miyoshi et al., 2010). In addition, the MGE generates prototypic neurons of the globus pallidus (Dodson et al., 2015), and basal forebrain cholinergic neurons (Allaway and Machold, 2017), while the CGE contributes to numerous amygdala nuclei (Tang et al., 2012). The LGE generates direct and indirect spiny projection neurons (MSNs) of the striatum (Yun et al., 2003), arkypallidal neurons of the globus pallidus (Dodson et al., 2015), olfactory bulb (OB) interneurons (Yun et al., 2003), as well as neurons of the olfactory tubercle and amygdala (Ko et al., 2013).

Several transcription factors (TFs) and their co-factors have been shown to be necessary for the specification of GABAergic subtypes (Leung et al., 2022; Flames et al., 2007), and their dysregulation results in disease (Leung et al., 2022; Zug, 2022). For example, members of the DLX family are present in the GE and are required for the development of GABAergic neurons (Anderson et al., 1997; Stühmer et al., 2002; Lindtner et al., 2019). The LIM homeodomain TF LHX6 is one of the factors known to regulate the generation of MGE-derived INs (Sandberg et al., 2016; Zhao et al., 2008), whereas MEIS2, a member of the TALE family of homeodomain-containing TFs, has been implicated in the generation of LGE-derived GABAergic PNs (Su et al., 2022). Haploinsufficiency of *MEIS2* in humans results in cardiac and palate abnormalities, developmental delay, and intellectual disability (Louw et al., 2015; Douglas et al., 2018; Giliberti et al., 2020; Zhang et al., 2021). The mechanisms by which these TFs select and activate their targets remain unclear.

Here, we used sparse CRISPR/Cas-mediated perturbation of *Meis2*, *Lhx6* and *Tcf4* in GABAergic progenitors and tracked their developmental trajectories with lineage barcodes and single-cell RNA sequencing (scRNA-seq). We found that the sparse perturbation of *Meis2* in the GE alters the development of GABAergic neurons, increasing the proportion of IN clones at the expense of PN clones. We identified genomic binding sites of MEIS1/2 in enhancers of genes that are differentially expressed in GABAergic PNs. MEIS1/2’s binding sites frequently overlapped with binding sites of DLX5 and LHX6. We performed luciferase reporter assays and found that only in the presence of DLX5 was MEIS2 able to activate the enhancers of PN genes. LHX6 repressed this DLX5/MEIS2-induced cooperative activation of PN genes, thus likely promoting an IN fate. Finally, a mutation of *Meis2* that causes intellectual disability in humans (Giliberti et al., 2020; Gangfuß et al., 2021) was much less able to potentiate the DLX5-induced activation of these enhancers. Our results indicate that MEIS2 acts as a transcriptional activator to generate patterns of enhancer activation that specifies PN identities within GABAergic precursor cells. This mechanism may contribute to neurological dysfunction in diseases caused by *MEIS2* mutations.

## Results

### *In vivo* tCROP-seq to assess the function of MEIS2 during fate decisions in GABAergic precursors

We conducted a logistic regression analysis on scRNA-seq data from the GE (Bandler et al., 2022) to identify regulatory TFs that play a role in determining the fate of GABAergic PNs or INs. Our findings revealed *Meis2* as the gene with the highest predictability for a PN fate, while *Lhx6* and *Tcf4* emerged as strong predictors of an IN fate (Figure 1a, S1a). To investigate the effects of *Meis2* perturbation on cellular fate decisions in a sparse population of precursors in the GE, we modified CROP-seq (Datlinger et al., 2017), a powerful method that enables pooled CRISPR screens while simultaneously capturing the transcriptome of individual cells. In our study, we focused on a sparse population of precursors within the GE to investigate the impact of the depletion of *Meis2* on fate decisions. Instead of lentiviral vectors to deliver single-guide RNAs (sgRNAs), our modified approach used a PiggyBac **t**ransposon-based strategy (**t**CROP-seq) and *in utero* electroporation to efficiently deliver sgRNAs to cycling progenitors in the GE (Figure 1b). The transposon system allows genes to be stably integrated into the genomes of electroporated cells and thus to be transmitted to their postmitotic daughter cells (Ding et al., 2005). This increases the pool of perturbed cells and ensures that the perturbation occurs during a period covering the peak of neurogenesis (Bandler et al., 2022). We also added specific capture sequences to the sgRNA vectors that efficiently link sgRNAs to cell barcodes, and enable sequencing of the protospacer from the transcriptome (Replogle et al., 2020). tCROP-seq sgRNA vectors also encode TdTomato to enable the labeling and enrichment of perturbed neurons. The efficiency of sgRNA *Meis2* to induce frame-shift mutations was validated *in vitro* and *in vivo* prior to the tCROP-seq experiments (Table S1).

**Figure 1:**
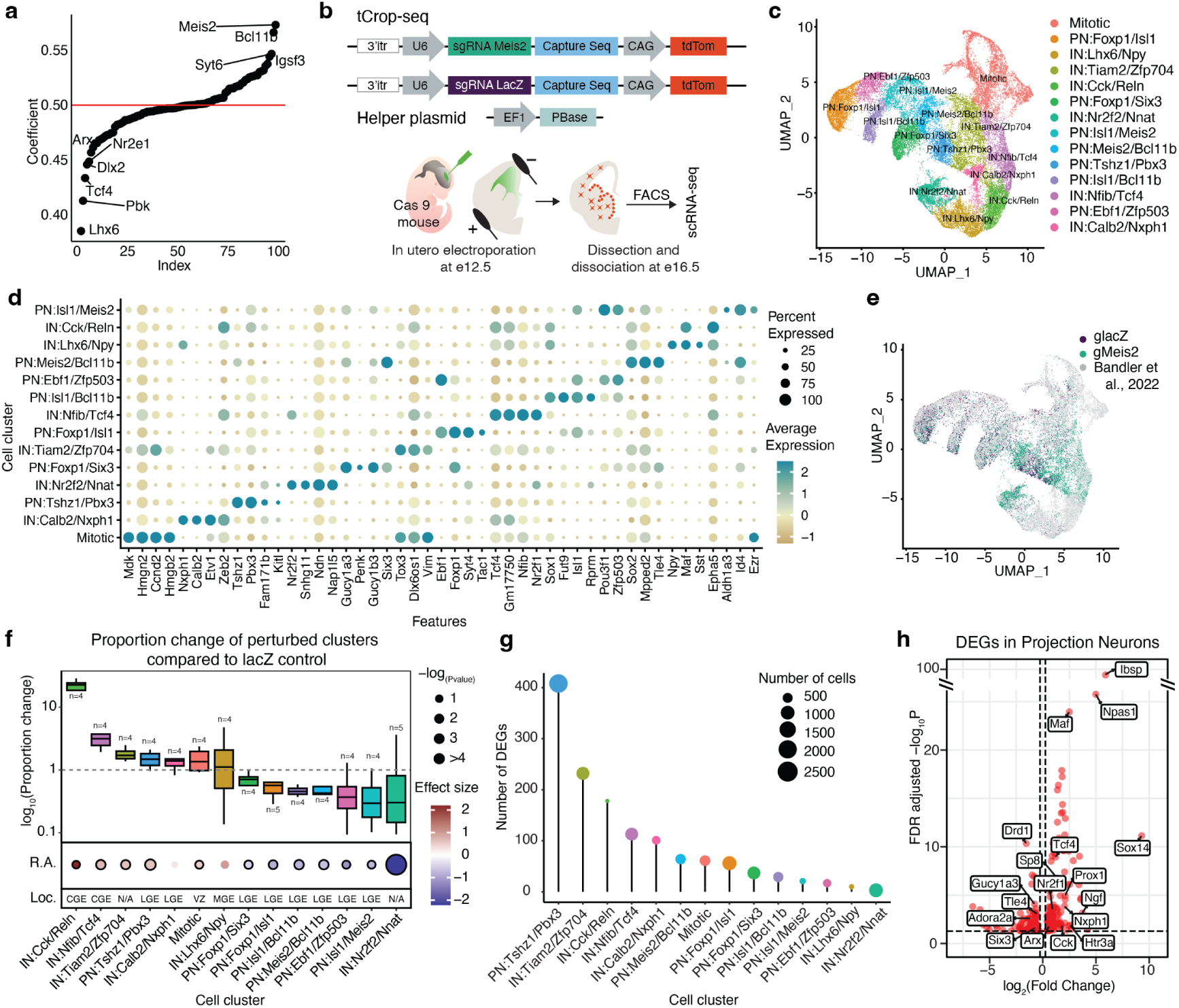
*In vivo* tCROP-seq of *Meis2* in the mouse forebrain. **a**, Logistic regression coefficients of genes being predictive for interneuron or projection neuron fate. Genes with coefficients >0.5 are predictive of projection neuron fate, and genes with coefficients <0.5 are predictive of interneuron fate. **b**, Vector maps and schematic of the *in vivo* tCROP-seq workflow, in which mutations in individual genes are introduced *in utero* and the effect is determined at a later time point via scRNA-seq. **c**, Uniform Manifold Approximation and Projection (UMAP) plot of inhibitory cells colored by clusters. **d**, Dotplot of the top four marker genes of inhibitory clusters. **e**, UMAP plot of the integrated dataset colored by sgRNAs. Grey dots represent cells from a published dataset (Bandler et al., 2022). **f**, Top: Relative increase or decrease in the number of inhibitory cell clusters in gMeis2 compared to gLacZ. Bottom: Perturbation effects in different clusters compared to lacZ controls. Dot color corresponds to effect size, dot size corresponds to negative base 10 log(P-value). P-values were calculated from linear modeling, Padj was calculated by Benjamini & Hochberg FDR correction. The black outline indicates statistical significance (p-val < 0.05). R.E., Regression Analysis; Loc., Location of the presumed origin of the cluster within the GE. **g**, Lollipop plots showing the number of differentially expressed genes (DEGs) for gMeis2 in inhibitory clusters. **h**, Volcano plot depicting differentially expressed genes in gMeis2 and gLacZ projection neurons.

The tCROP-seq vectors were targeted by *in utero* electroporation at E12.5 to progenitor cells of the GE in a mouse line ubiquitously expressing Cas9 (Platt et al., 2014) (Figure 1b). At E16.5, most TdTomato+ cells had migrated away from the ventricular zone (VZ) and colonized a variety of structures, including the striatum, cerebral cortex, and OB (Figure S1b-c), consistent with the migration patterns of GE-derived inhibitory neurons at this stage (Anderson et al., 2001). Both immunohistochemical analysis of TdTomato+ cells at E18 and scRNA-seq analysis at E16 showed that the tCROP-seq vectors were expressed across a variety of MGC, CGE and LGE derived inhibitory neuron types (Figure S1b, see below).

For the tCROP-seq experiment, we collected a total of 14 embryos from 10 pregnant females (Table S9). Of these, 8 received sgRNAs for *Meis2* (gMeis2) and 6 received sgRNAs for *LacZ* (gLacZ), which served as a control. Cortices, striata, and OBs were dissected at E16 and TdTomato+ cells were enriched by FACS. tCROP-seq allows the retrospective assessment of which sgRNA was expressed in which cell. We pooled cells from embryos having received gLacZ or gMeis2, and conducted multiplexed single-cell RNA sequencing to minimize batch effects (Figure 1b; see Methods) (Jin et al., 2020). We sequenced 6 independent scRNA-seq experiments. Together, this resulted in a dataset containing 34481 cells passing quality controls and filtering, that were linked with either gLacZ (11009 cells) or gMeis2 (23472 cells). We projected cells into a shared embedding using Harmony (Korsunsky et al., 2019) and applied a standard Seurat (Hao et al., 2021) analysis pipeline (Figure S1d).

### Single-cell perturbation of Meis2 alters the proportion of PNs and INs

Louvain clustering grouped glia cells, excitatory neurons, and inhibitory neurons into multiple clusters (Figure S1d). We subset cells from inhibitory clusters (16098 inhibitory cells; Figure S1e-h) and integrated them with published scRNA-seq datasets from embryonic wild-type mice (Bandler et al., 2022), to get a higher resolution of inhibitory cell states (Figure 1c). We annotated 14 inhibitory clusters based on shared marker gene expression and grouped them into three major classes: mitotic (mitotic), GABAergic PNs (PN:Foxp1/Six3, PN:Foxp1/Isl1, PN:Isl1/Bcl11b, PN:Ebf1/Zfp503, PN:Meis2/Bcl11b, PN:Isl1/Meis2, PN:Tshz1/Pbx3), and GABAergic INs (IN:Calb2/Nxph1, IN:Tiam2/Zfp704, IN:Nfib/Tcf4, IN:Lhx6/Npy, IN:Cck/Reln, IN:Nr2f2/Nnat; Figure 1c-d, Table 1, Table S2). Cells expressing gMeis2 contained a reduced proportion of PN cell-types and an increased proportion of IN cell-types, when compared to gLacZ controls (Figure 1f). Interestingly, the proportion of CGE-derived IN populations was increased in the gMeis2 condition, and the relative proportion of multiple PN types was decreased. This suggests that, under normal conditions, MEIS2 promotes the generation of LGE-derived PN types at the expense of CGE-derived IN types. A pseudo-bulk differential gene expression analysis (DEG) (Squair et al., 2021) of GABAergic neurons comparing gMeis2 and gLacZ showed reduced expression levels of genes known to be involved in PN development and increased expression levels of genes known to be involved in IN development (Table S3). The impact of gMeis2 on differential gene expression was strongest on the clusters PN:Tshz1/Pbx3, IN:Tiam2/Zfp704 and IN:Cck/Reln (Figure 1g, S2a, Table S4). In PN clusters, gMeis2+ cells showed decreased expression levels of genes known to be associated with PN identity, such as *Adora2a*, *Drd1*, and *Six3* (Kreitzer and Malenka, 2008; Song et al., 2021; Knowles et al., 2021), compared to gLacZ (Figure 1h, Table S3-4). Many genes related to IN development and specification, such as *Maf*, *Tcf4*, *Prox1*, *Arx* (Lim et al., 2018; Miyoshi et al., 2015; Batista-Brito et al., 2008), were up-regulated in PN clusters (Figure 1h, Table S3-4). Futhermore, also the proportion of mitotic progenitors was increased in gMeis2 compared to gLacZ. Genes involved in cell proliferation and differentiation were up-regulated in the mitotic cluster in gMeis2, in particular the gene *Wnt5a*, which is part of the non-canonical WNT signalling pathway (Chenn and Walsh, 2002; Megason and McMahon, 2002) (Figure 1f, S2b,c). GO Term analysis of the up and down-regulated DEGs revealed that processes, such as neuron development, axon extension, and neuron differentiation, were deregulated (Figure S2d).

**Table 1:** GABAergic precursor clusters and associated brain regions. The table presents E16 clusters of GABAergic neuronal precursors along with their corresponding descriptions and associated brain regions. At E16, these scRNA-seq clusters represent precursors of adult neuronal types, many of which are in the process of migration to their final settling positions. Due to the ongoing migration and developmental processes, the specific type they will differentiate into and the structure they will migrate to can only be inferred or hypothesized (Mayer et al., 2018; Lee et al., 2022; Lim et al., 2018; Bandler et al., 2022). We have inferred these potential future fates based on Mousebrain.org (La Manno et al., 2021) and DropViz.org (Saunders et al., 2018). AMY, Amygdala; BNST, Bed nucleus of the stria terminalis; CGE, caudal ganglionic eminence; MGE, medial ganglionic eminence CTX, Cortex; GP, Globus pallidus; HC, Hippocampus; ITC, intercalated cells; MSN, Medium spiny neuron; D1-MSN, DRD1-expressing MSN (direct pathway striatal projecting neuron); D2-MSN, DRD2-expressing MSN (indirect pathway striatal projecting neuron); Pvalb, Parvalbumin expressing interneuron; OB, Olfactory bulb; Sst, Somatostatin expressing interneuron; Th INs, TH expressing interneuron; Reelin, Reelin expressing interneurons; VIP, VIP expressing interneuron;

| Cluster | Description | Region |
| --- | --- | --- |
| Mitotic | Mitotic cells based on high expression of cell-cycle related genes | VZ |
| IN:Calb2/Nxph1 | Precursor of Sst, Pvalb and Th INs | STR, CTX, OB |
| PN:Tshz1/Pbx3 | Precursor of MSNs, ITCs | BNST, AMY |
| IN:Nr2f2/Nnat | Precursor of CGE derived INs | CTX, HC |
| PN:Foxp1/Six3 | Precursor of D2-MSN, Ppp1r1b-type PN | STR, GP |
| IN:Tiam2/Zfp704 | Precursor of CGE derived INs: Sst, Pvalb INs | CTX, HC, OB |
| PN:Foxp1/Isl1 | Precursor of MSN | STR |
| IN:Nfib/Tcf4 | Precursor of CGE derived INs | CTX, HC, OB |
| PN:Isl1/Bcl11b | Precursor of D1-MSN and Pp1r1b-type PN | STR, GP |
| PN:Ebf1/Zfp503 | Precursor of D1-MSN | STR |
| PN:Meis2/Bcl11b | Precursor of MSN | STR, AMY |
| IN:Lhx6/Npy | Precursor of MGE derived INs: Pvalb, Sst | CTX, HC |
| IN:Cck/Reln | Precursor of CGE derived INs: Vip, Reelin | CTX, HC |
| IN:PN:Isl1/Meis2 | Precursor of MSN | AMY, BNST |

### Combined *in vivo* lineage tracing and tCROP-seq reveals a shift in clonal compositions of perturbed cells

Our findings thus far raised the question of how neurons with a broad PN identity (Louvain clustering grouped them into PNs) acquired CGE/MGE-IN signatures. One possibility would be that the perturbation in gMeis2 alters the cell cycle dynamics of PN progenitors or that PN progenitors undergo cell death. Alternatively, progenitors of the LGE-PN lineage may fail to establish a proper PN identity and switch to a CGE/MGE-IN identity. To test these possibilities, we combined tCROP-seq with a barcode lineage tracing method called TrackerSeq (Bandler et al., 2022)), that integrates heritable DNA barcodes into the genome of electroporated progenitors, enabling the tracking of clonal relationships between their daughter neurons (Figure 2a). tCROP-seq and TrackerSeq can be used simultaneously because we have implemented a similar transposase strategy for both methods (Figure 2a). We used *in utero* electroporation at E12.5 to introduce the TrackerSeq barcode library and tCROP-seq sgRNAs to cycling progenitors in the GE. We collected TdTomato/EGFP+ cells from 4 independent batches and prepared sequencing libraries for transcriptomes, sgRNAs, and lineage barcodes. The cells with TrackerSeq barcodes were part of the preceding tCROP-seq analysis and were thus integrated in the same embedding (Figure 2b). Consistent with Bandler *et al*. 2022 (Bandler et al., 2022), we found clones composed of mitotic cells, PNs, INs, and combinations thereof (Figure 2c-d). The average clonal size of multi-cell in gMeis2 was unchanged compared to gLacZ (Figure 2e and Figure S2e-f), suggesting that cell cycle dynamics or cell-death are unlikely to be responsible for the observed proportional shift in cell fate. The proportion of clones consisting of only mitotic cells was relatively increased in gMeis2 compared to gLacZ, which agrees with a report showing that MEIS2 is required for LGE progenitors to leave the cell cycle (Su et al., 2022) (Figure 2f). 225 clones dispersed across mitotic and PN clusters (mitotic-PN), and 100 clones dispersed across mitotic and IN clusters (mitotic-IN; Figure 2f). Strikingly, when we compared clonal patterns of gMeis2 and gLacZ cells, we observed a pronounced shift toward IN-only and mitotic-IN clones. Conversely, the number of PN-only, and mitotic-PN clones was decreased (Figure 2f). Furthermore, the coupling of multi-cell clones within interneuron clusters was reduced in gMeis2, which may indicate that a wider range of lineages develop into interneuron precursors as a result of the fate switch from PNs to INs (Figure S2g). Our results indicate that perturbation of progenitors with gMeis2 leads to a partial change in the fate of newly generated neurons, resulting in a bias in favour of INs instead of PNs.

**Figure 2:**
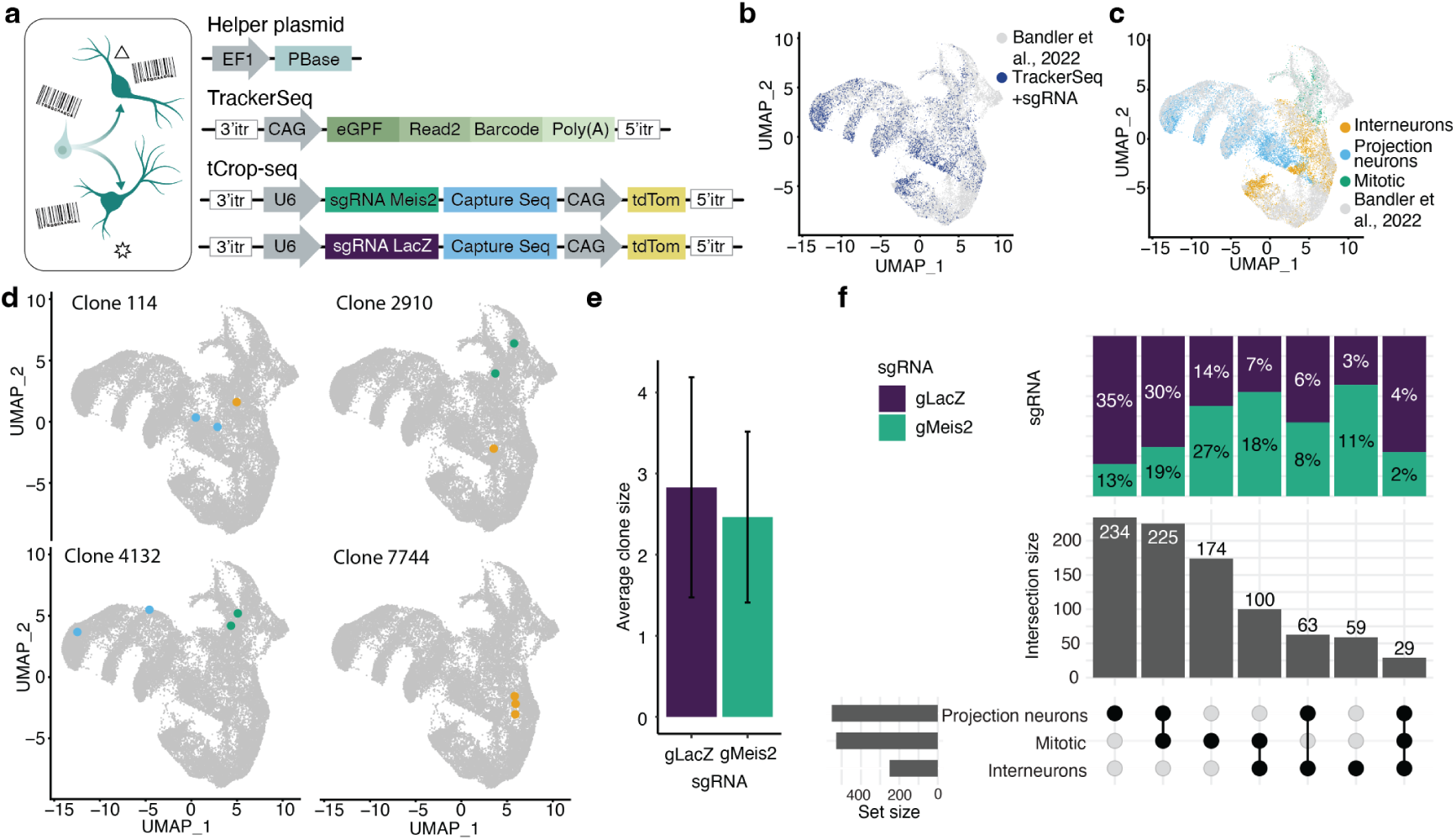
*In vivo* TrackerSeq lineage tracing and tCROP-seq perturbation of *Meis2*. **a**, Schematic of TrackerSeq lineage tracing, in which clonal boundaries are determined using heritable RNA tags. **b**, UMAP of the integrated dataset with labelling of cells containing TrackerSeq lineage barcodes. **c**, UMAP of the integrated dataset colored by cell class (mitotic, interneurons, projection neurons). **d**, Examples of clones that are shared between classes, and an example of a clone restricted to one class. **e**, Bar graph depicting the average clone size of inhibitory clones in the gLacZ and gMeis2 datasets. **f**, UpSet plot showing clonal intersections between cell classes. The bar graph on top displays the proportion of clones belonging to gLacZ or gMeis2. The bar graph in the middle shows the number of observed intersections. The bar graph on the left indicates the number of cells per cluster.

### Genomic binding of DLX5 and MEIS2 in the embryonic GE

To identify target genes of MEIS2, we performed chromatin immunoprecipitation-sequencing (ChIP-seq) on GE tissue dissected from E14.5 mouse embryos, using a combination of anti-MEIS1/2 and anti-MEIS2 antibodies. In the GE, the expression of *Meis2* is higher and more widespread than that of *Meis1*, therefore the antibodies are likely to bind primarily to MEIS2 epitopes (Figure S3a-b). We identified 3780 MEIS1/2 binding sites, of which 16% were located within 5 kb of a transcription start site (TSS; Figure 3a). 20% of the biding sites overlapped with developmental enhancers linked to putative target genes (Gorkin et al., 2020), Table S5). Our data predict that MEIS1/2 directly regulates 1218 target genes, either by binding to their TSS or distal enhancers. Many of the target genes (16%) overlapped with genes that were up-regulated in gMeis2-tCROP-seq positive PNs cells (Figure 3b, Table S4-5). *De-novo* motif analysis revealed the previously described MEIS1/2 core hexameric and decameric binding motifs TGACAG and TGATTGACAG, which were highly enriched at the centers of the peaks. These motifs correspond to either the binding of the MEIS homodimer, or the MEIS/PBX heterodimer, respectively (Chang et al., 1997; Shen et al., 1997) (Figure 3c, S3b). Binding motifs containing the core sequence TAATT were strongly enriched in MEIS1/2 ChIP-seq peaks, and enriched at enhancers compared to TSS-associated regions. This motif is shared by several homeodomain TF families including those of DLX, LHX and ISL (Figure 3d) (Leung et al., 2022), of which several members are expressed in the GE (Mayer et al., 2018; Leung et al., 2022; Flames et al., 2007). Among them, we found the strongest enrichment for the binding motif of DLX3 (Figure 3d).

**Figure 3:**
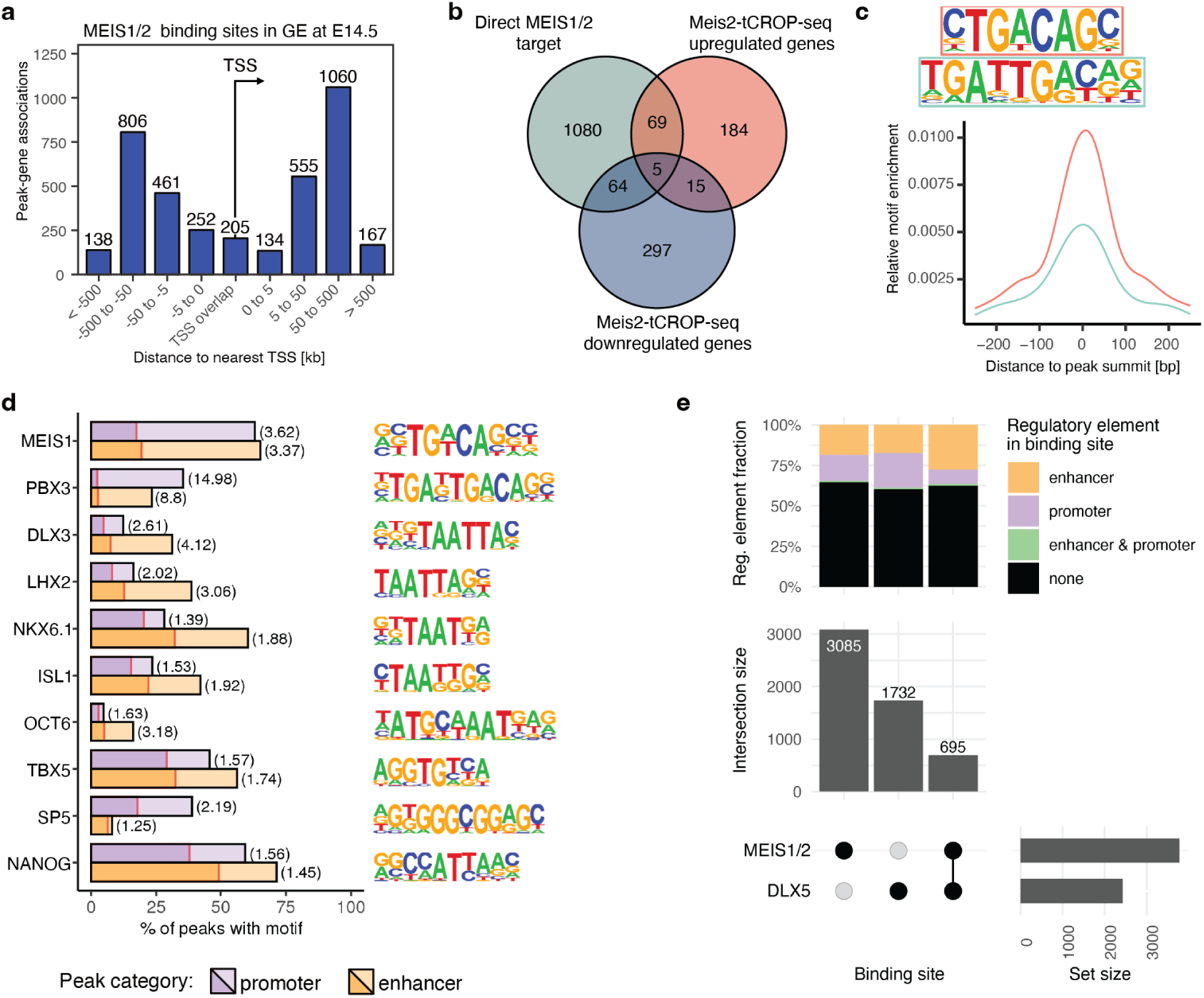
DNA binding sites of MEIS1/2 in the GE at E14.5. **a**, Distribution of MEIS1/2-ChIP-seq peaks relative to the nearest transcriptional start site (TSS). **b**, Venn diagram showing overlap between MEIS1/2 target genes and genes up- or downregulated in inhibitory neurons of gMeis2-tCROP-seq (*p*_*val*_*ad* *j*_ < 0.05&(*avg*_*logFC* < −1.0|*avg*_*logFC* > 1.0)). Overlap of up- and down-regulated genes is due to opposite regulation in different subtypes of inhibitory neurons. **c**, *De novo* identified MEIS1/2 binding motifs and their position relative to peak summits. **d**, Motif occurrence of selected known motifs enriched within enhancer- or promoter-overlapping MEIS1/2 binding sites (light bars) compared to G/C-matched reference sequences (dark bars), with fold-enrichment in parentheses. **e**, Overlap between binding sites of MEIS1/2 and DLX5 (bottom), with respective distribution of binding sites overlapping promoter and/or enhancer regions.

All DLX TFs share a common conserved motif, of which DLX1, DLX2, DLX5, and DLX6 are known to be master regulators of inhibitory neuron development in the forebrain (Lindtner et al., 2019; Panganiban and Rubenstein, 2002). Because *Meis2* and *Dlx5* are co-expressed in PN precursor cells of the GE (Figure S4g, S6), we next tested if MEIS2 and DLX5 interact in the GE. First, we compared the binding sites of MEIS1/2 with those of a published DLX5 ChIP-seq dataset in mouse GE (Lindtner et al., 2019). Numerous MEIS1/2 binding sites (695; 18%) overlapped with DLX5 binding sites. Remarkably, the proportion of enhancers at shared (MEIS1/2-DLX5) binding sites was significantly increased compared to MEIS1/2- and DLX5-exclusive binding sites (Figure 3e; p = 8.856e-9, Chi2-test). The spacing and orientation of MEIS and DLX motifs have previously been described *in vitro*, and changes in spacing between co-transcription factors have been shown to affect gene regulatory capacity (Jolma et al., 2015; Ng et al., 2014; Jindal and Farley, 2021). In our data, the most common motif spacing was 2-4 bp. In contrast to published *in vitro* experiments that observed a fixed spacing of 2 bp between MEIS1 and DLX3 (Jolma et al., 2015), we observed a wider range of spacing (Figure S3d). Together, our findings suggest a potential cooperative role of MEIS1/2 and DLX5 in the fate determination of GE-derived neurons.

### Functional link between MEIS2/DLX5 and PN fate

To investigate the possibility of a functional link between MEIS2 and DLX5 in PN development, we performed a series of dual luciferase reporter assays to measure the activity of select genomic enhancers in the presence of MEIS2, DLX5, or both. To select enhancers active in the developing forebrain, we intersected MEIS1/2-DLX5 co-binding sites from ChIP-seq data with the VISTA *in vivo* enhancer database (Visel et al., 2007) (Figure S3e). Additionally, we confirmed the accessibility of the respective genomic regions, utilizing published scATAC-seq data of the LGE and MGE (Rhodes et al., 2022) (Figure 4a). First, we chose two enhancers (hs1080 and hs956) of the TF *Foxp2*, which both contained MEIS/DLX motifs with a spacing of 3 bps (Visel et al., 2007; Visel et al., 2013) (Figure 4a, Figure S4a, b, d, e). *Foxp2* is expressed in precursors of GABAergic PNs (Figure S4g), has previously been implicated in PN development (den Hoed et al., 2021; French and Fisher, 2014), and is one of the genes that we found to be downregulated in gMeis2 tCROP-seq experiments (Table S4). We transfected Neuro2a cells with a plasmid containing a selected enhancer upstream of a minimal promoter and the firefly luciferase gene, as well as a control plasmid encoding the NanoLuc luciferase gene under the PGK promoter. Additionally, we transfected the cells with plasmids encoding *Dlx5*, *Meis2*, or both. If the enhancer can be activated by DLX5, MEIS2, or both, the transfected cells would produce measurable luciferase activity. MEIS2 alone did not significantly activate either enhancer, and both *Foxp2* enhancers were only modestly activated in the presence of DLX5 alone (Figure 4b-c). Remarkably, MEIS2 and DLX5 together strongly potentiated the DLX5-induced activation of the *Foxp2* enhancers. As expected, PBX1, a known interaction partner of MEIS2 (Hyman-Walsh et al., 2010), increased the effect of MEIS2 (Figure S4c,f). These results suggest that MEIS2 and DLX5 bind cooperatively at specific binding sites of enhancers to regulate *Foxp2* expression.

**Figure 4:**
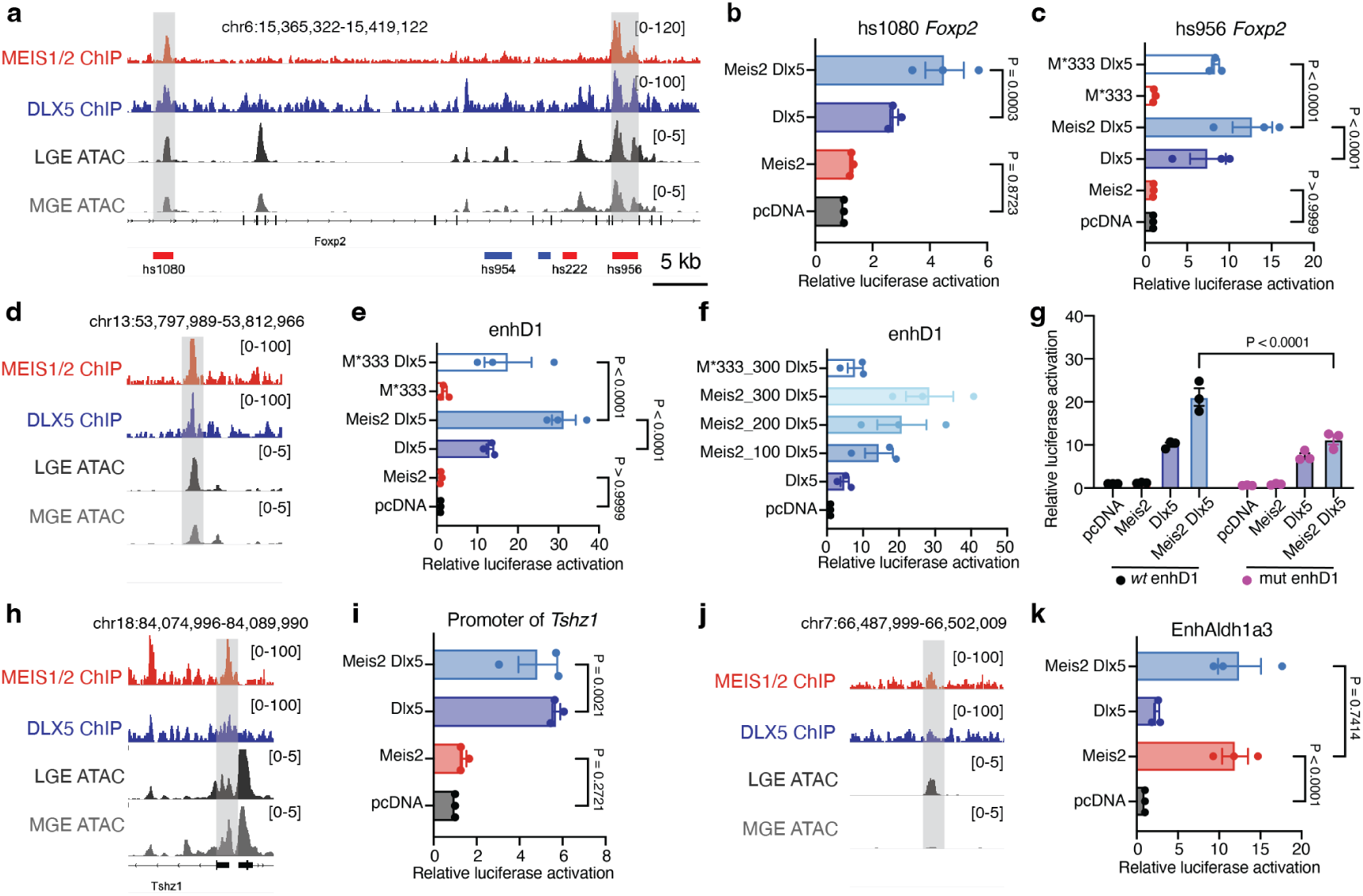
Cooperation between MEIS2 and DLX5 activates enhancers of projection neuron-specific genes. **a**, Representative profiles of MEIS1/2 (red) and DLX5 (blue) ChIP-seq at E14.5 and E13.5 respectively, as well as scATAC-seq from LGE (dark gray) and MGE (gray) at E12.5 are shown at the *Foxp2* gene locus. DLX5 ChIP-seq data from (Lindtner et al., 2019); scATAC-seq data from (Rhodes et al., 2022). **b**, Luciferase activity driven by the enhancer hs1080, co-transfected with *Meis2* and *Dlx5* expression vectors in Neuro2a cells. **c**, Luciferase reporter assays of the enhancer hs956. **d**, Representative profiles of the *Drd1* gene enhancer enhD1. **e**, Luciferase reporter assays of enhD1. **f**, Luciferase reporter assays of enhD1, co-transfected with Dlx5 and increasing concentration of Meis2, or with Meis2*333. **g**, Luciferase reporter assays of the wild-type or mutated, shorter version of enhD1. **h**, Representative profiles of the *Tshz1* promoter. **i**, Luciferase reporter assays of the *Tshz1* promoter. **j**, Representative profiles of the *Aldh1a3* enhancer enhAldh1a3. **k**, Luciferase reporter assays of enhAldh1a3. In panels b, c, e, f, g, i and k, bars represent mean ± s.e.m from a total of 9 replicates, split into three independent batches, each performed in triplicate. Points represent the mean of each batch for each condition. Statistical significance was assessed by two-way ANOVA. P-values of pairwise comparisons from post-hoc Tukey’s HSD are presented for selected conditions.

Mutations in the *MEIS2* gene have been linked to intellectual disability, cardiac defects and facial phenotypes (Louw et al., 2015; Verheije et al., 2019; Giliberti et al., 2020; Gangfuß et al., 2021). At least four patients with severe disease carry either a frameshift mutation, an in-frame deletion, or a missense mutation of a single highly conserved amino acid (Arg333) located in the MEIS2 homeodomain (Giliberti et al., 2020; Gangfuß et al., 2021). We tested whether the p.Arg333Lys missense variant (MEIS2*333) can activate the *Foxp2* enhancer hs956. DLX5-dependent joint activation of hs956 was greatly reduced with MEIS2*333 compared to wild-type MEIS2 (Figure 4c).

Next, we investigated whether the cooperation of MEIS2 and DLX5 at co-binding sites activates a putative regulatory enhancer (enhD1) of *Drd1*. *Drd1* encodes for the dopamine receptor D1, which is a top marker of D1-type medium spiny projection neurons (D1-MSN; PN:Foxp1/Isl1, PN:Isl1/Bcl11b, PN:Ebf1/Zfp503) in the striatum (Gerfen and Surmeier, 2011) (Figure 4d, Figure S4g). Its gene expression was strongly reduced in PN clusters in gMeis2 tCROP-seq experiments (Table S4, Figure 1h). EnhD1 is predicted to be associated with *Drd1* (Figure S4h) (Gorkin et al., 2020) and is located in the same topologically associated domain (TAD) (Bonev et al., 2017). Furthermore, enhD1 contained pronounced ChIP-seq peaks for DLX and MEIS1/2 (Figure 4d), and multiple MEIS/DLX co-binding motifs (Figure S4i). Similar to the *Foxp2* enhancers, MEIS2 did not activate enhD1, but it potentiated the effect of DLX5 on the activity of enhD1, in a concentration-dependent manner (Figure 4e-f). The cooperative activation of enhD1 by MEIS2 and DLX5 was greatly reduced with the mutated version of MEIS2 (MEIS2*333; Figure 4f). A truncated version of enhD1 in which a portion (TG) of the MEIS binding motif was removed at multiple sites of the enhancer (Figure S4i), showed reduced activation by MEIS2/DLX5 compared with the unmodified truncated enhD1 (Figure 4g). Taken together, our findings suggest that the cooperation of MEIS2 and DLX5 at specific co-binding sites within *cis*-regulatory elements activates PN-specific gene expression to promote PN fate.

Next, we tested whether MEIS2 is able to activate the promoters of its target genes *Pbx3*, *Tshz1*, *Zfp503*, and *Six3*. All three genes are marker genes for different PN clusters, and they all contain binding sites for MEIS in their promoters (Figure 1c, S5a-c). We found that the activation of these promoters by MEIS2 is small (Figure S4h). Interestingly, even the *Tshz1* promoter, which contains both DLX5 and MEIS1/2 motifs, was not activated by MEIS2, nor was MEIS2 able to enhance the DLX5-induced activation of this promoter (Figure 4h-i). This may be because the motifs for MEIS1/2 are far away from DLXs motifs (Figure 3d).

Our data suggest that in the GE, MEIS2 requires the presence of DLX5 to bind and co-activate *cis*-regulatory enhancers with specific co-binding sites, and this process induces gene expression related to PN development. We performed additional luciferase reporter assay experiments where we included Dlx1, Dlx2, Dlx6 and expanded the analysis of the ChIP-seq datasets to include additional members of the Dlx family (Dlx1, Dlx2). Our data indicate that Dlx1/2/6 TFs play a similar role as Dlx5, and can activate the tested enhancers (Figure S11i-k, Table S5).

We tested a total of 8 enhancers of genes which are known to be important for inhibitory neuron development using a dual-luciferase reporter assay (Figure S5g-h), and the results support this model. Of the enhancers tested, only the LGE-specific enhancer of *Aldh1a3*, enhAldh1a3, which lacks a MEIS1/2-DLX5 co-binding site, was strongly activated by MEIS2 alone (Figure 4j-k; Figure S4g). *Aldh1a3* encodes an enzyme that synthesizes retinoic acid in LGE precursors at E12.5 (Molotkova et al., 2007; Toresson et al., 1999) and is essential for the differentiation of striatal PNs (Chatzi et al., 2011). *Aldh1a3* was greatly downregulated in several clusters in the gMeis2 tCROP-seq experiments (Table S4). It remains unclear whether MEIS2 is able to activate enhAldh1a3 on its own, or whether another co-factor, present in Neuro2a cells, is required.

### Spatial patterning and the functional activity of MEIS2 in the GE

PNs of the striatum originate largely in the LGE, and many IN types, *e.g.*, those of the cortex, originate in the MGE and CGE (Knowles et al., 2021; Lim et al., 2018; Bandler et al., 2017). *Meis2* mRNA is initially expressed broadly in the VZ of the LGE, CGE and MGE. In neuronal precursors of the subventricular (SVZ) and mantle zones (MZ), a spatial pattern of *Meis2* expression emerges, where *Meis2* continues to be highly expressed in the LGE, but is absent in the MGE (Figure S6) (Toresson et al., 1999; Su et al., 2022).

We next asked how the function of MEIS2 as a DLX-dependent activator of PN development acquires LGE selectivity. We argued that LHX6 might be involved in this process. First, the mRNA expression pattern of *Lhx6* contrasts with that of *Meis2*, being exclusively expressed in the MGE and enriched in the SVZ and MZ (Figure S6) (Flames et al., 2007). Consistently, we found only a small population of cells at the interface of the VZ and SVZ in the MGE, that showed co-immunoreactivity of MEIS2 and LHX6 (Figure 5a). Second, LHX6 is a strong predictor of IN fate (Figure 1a) and its activity is known to be required for the specification of cortical IN subtypes (Sandberg et al., 2016; Zhang et al., 2013; Cesario et al., 2015).

**Figure 5:**
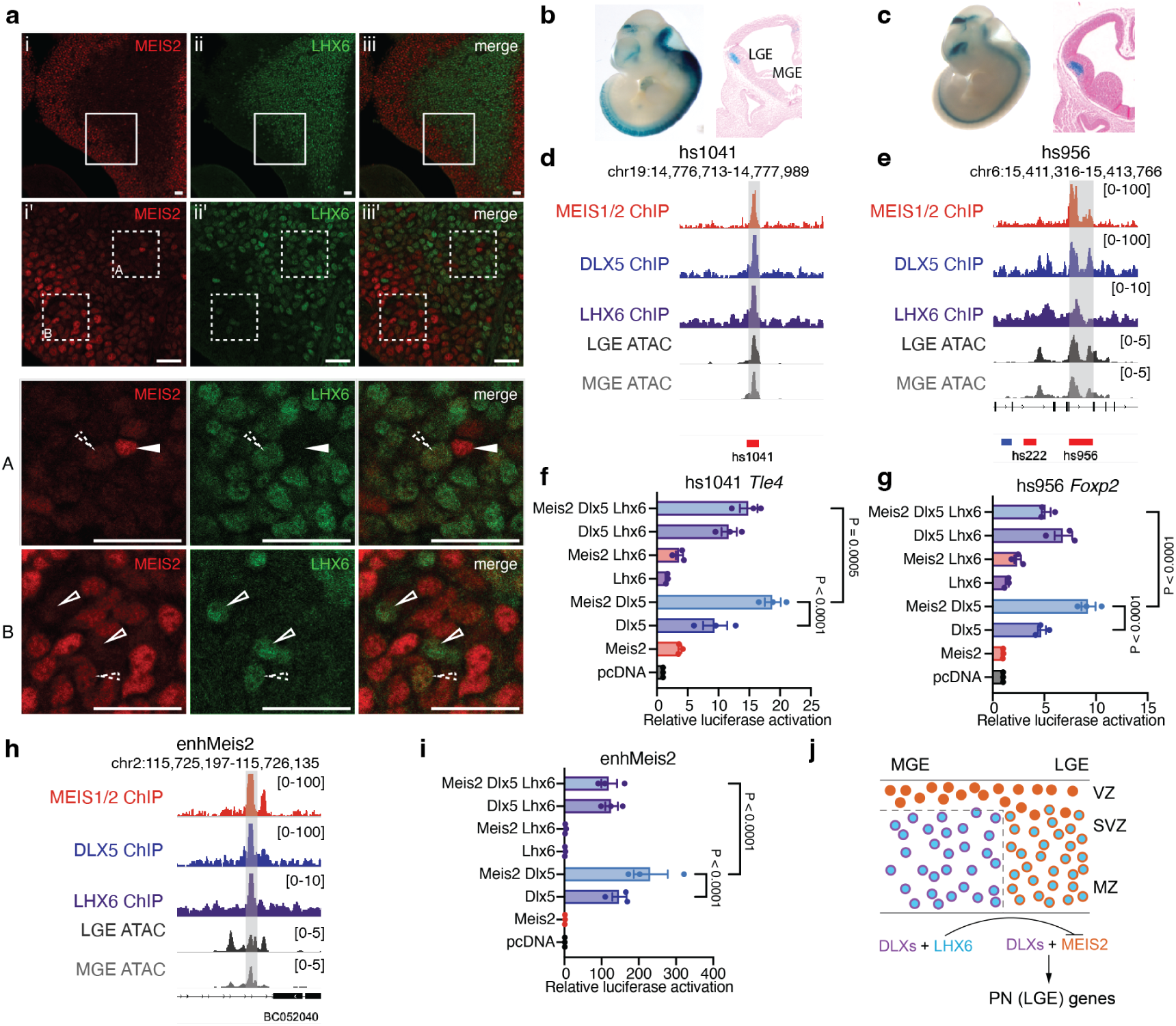
Regulation of LGE enhancers by MEIS2, DLX5 and LHX6. **a**, Immunohistochemistry of MEIS2 and LHX6 in the MGE of E13.5 embryos. MEIS2 immunoreactivity is high in cells of the VZ and low as cells transition to the VZ/MZ. Few cells in the SVZ retain MEIS2 expression (white triangle). Conversely, few cells in the VZ are immunoreactive for LHX6 (empty triangles). Some cells at the VZ/SVZ interface are co-immunoreactive against MEIS2 and LHX6 (dotted triangles). **b-c**, LacZ expression in the LGE of E12.5 embryos driven by the enhancers hs1041 and hs956 (Visel et al., 2007). **d-f**, Representative tracks of MEIS1/2 ChIP-seq in the GE at E14.5 (red), DLX5 ChIP-seq in the GE at E13.5 (blue) (Lindtner et al., 2019), LHX6 ChIP-seq in the GE at E13.5 (purple) (Sandberg et al., 2016) and scATAC-seq in LGE (dark gray) and MGE (gray) at E12.5 (Rhodes et al., 2022). **g-i**, Luciferase activity driven by hs1041, hs956, and hs748 enhancers co-transfected with *Meis2*, *Dlx5*, and *Lhx6* expression vectors in Neuro2a cells. **j**, Representative tracks of enhancer enhMeis2. **k**, Luciferase reporter assays of enhMeis2. **l**, Model of the proposed actions of MEIS2, DLX5 and LHX6. MEIS2 promotes projection neuron fate in the presence of DLX. LHX6 represses Meis2 expression and function. SVZ, subventricular zone; VZ, ventricular zone; MZ, mantle zone. In panels g, h, i, f, bars represent mean ± s.e.m from a total of 9 replicates, split into three independent batches, each performed in triplicate. Points represent the mean of each batch for each condition. Statistical significance was assessed by two-way ANOVA. p-values of pairwise comparisons from post-hoc Tukey’s HSD are presented for selected conditions.

We intersected ChIP-seq peaks in the GE of MEIS1/2, DLX5 (Lindtner et al., 2019) and LHX6 (Sandberg et al., 2016) (Figure S3f, Table S5). Out of 151 MEIS1/2-DLX5-LHX6 overlapping peaks, 41 were within VISTA enhancers, and 28 of these enhancers showed activity in the developing forebrain (Figure S3g, S7). We selected three of them to perform luciferase reporter assays (Figure 5b-g, S3h-j): (1) hs1041, an enhancer of the *Tle4*, which encodes transcription co-repressor 4, (2) hs956, an enhancer of *Foxp2*, and (3) hs748, an enhancer of *Zfp503*, which encodes the zinc finger protein TF 503 (NOLZ1). Genes regulated by the selected enhancers are known to play a role in striatal development (Shang et al., 2022; den Hoed et al., 2021; French and Fisher, 2014; Su-Feher et al., 2021), were expressed in PN precursors (Table S2), and were reduced in several clusters in the gMeis2 tCROP-seq experiments (Table S4). Consistent with the above findings, MEIS2 strongly potentiated the DLX5-mediated activation of hs1041, hs956, and hs748 reporters. LHX6 alone had little to no effect on the activation of these enhancers. However, co-expression of LHX6 with MEIS2 and DLX5, resulted in a strong suppression of enhancer activity in all three cases (Figure 5f-g, S3j). This suggests that LHX6, whose expression is spatially restricted to the MGE, suppresses the DLX5-MEIS2-induced enhancer activation in the MGE. To gather further evidence for this mechanism, we screened 20 VISTA enhancers with overlapping ChIP-seq peaks for LHX6, MEIS1/2 and DLX5 (Figure S7). As expected, none of them exhibited robust activity in the MZ of the MGE. Next, we explored the putative enhancer of *Meis2*, enhMeis2 (Gorkin et al., 2020), which also contained MEIS1/2-DLX5-LHX6 co-binding sites (Figure 5h). MEIS2 strongly potentiated the DLX5-mediated activation of enhMeis2 (Figure 5i), suggesting that in the presence of DLX5, MEIS2 can promote its expression via the activation of enhMeis2. Self-activation has already been reported previously for *Meis* genes (Bridoux et al., 2020). Strikingly, LHX6 strongly repressed the MEIS2-DLX5 mediated activation of enhMeis2, suggesting that LHX6 suppresses the expression of MEIS2, consistent with a recent *Lhx6* knockout study in mice (Asgarian et al., 2022). This may explain the absence of MEIS2 in the SVZ and MZ of the MGE, and adds another level of regulation aimed at suppressing PN fate in MGE precursors (Figure 5j, S6). Together, LHX6 represses both MEIS2 gene expression and function in MGE.

### Meis2 and Lhx6 alter gene modules in PNs and INs

To explore how the depletion of embryonic TFs alters postnatal cell-type composition and identity, we performed pooled tCROP-seq experiments with sgRNAs for *Meis2* (gMeis2), *Lhx6* (gLhx6), *Tcf4* (gTcf4), and LacZ (gLacZ, control). Like LHX6, based on our regression analysis (Figure 1a) TCF4 is strongly predictive for an IN fate, but is expressed in all GE (Kim et al., 2020) (Figure S6). We delivered sgRNAs via *in utero* electroporation at E12.5 (Figure 6a-b), dissected 35 pups at P7, enriched TdTomato/EGFP positive cells with FACS, and performed pooled scRNA-seq. A total of ten scRNA-seq datasets were combined in silico, clustered, and annotated based on known marker genes (Figure 6c-d, S8, S9, Table S6, S9). All three perturbations had a significant effect on the composition of cell types compared to the gLacZ control (Figure 6e-f). As expected, cells expressing gLhx6 showed an increased proportion of medium spiny projection neurons (D1/D2 MSNs), OB precursors, and CGE INs compared to gLacZ. An increase of CGE INs after *Lhx6* deletion has previously been reported (Vogt et al., 2014). In addition, consistent with our embryonic tCROP-seq data, the proportion of INs was also increased in gMeis2 compared to gLacZ controls at P7. Furthermore, cells expressing gMeis2 showed a reduced proportion of intercalated cells of the amygdala (ITC), as well as OB inhibitory neurons and oligodendrocyte progenitors (Figure 6e-f). gTcf4 expression had a more modest effect on cell proportions, showing only a slight reduction in inhibitory neurons in the OB. Across all clusters, gLhx6, gMeis2, and gTcf4 positive cells had a total of 90, 58, and 7 DEGs respectively (Figure 6g-h, Table S7). Many of them were marker genes specifically expressed in IN or PN cell types (Table S6-7). gLhx6 perturbed cells were enriched for PN specific genes (*Isl1*, *Foxp1*, *Ebf1*, *Adora2a*, *Drd1*, *Six3*). By contrast, gMeis2 DEGs were enriched for IN-specific genes (*Maf* and *Prox1os*) and depleted for PN-specific genes (*Mpped2* and *Pbx3*). Our data support the conclusion that MEIS2 primarily induces PN fate and LHX6 primarily induces IN fate (Figure 1a).

**Figure 6:**
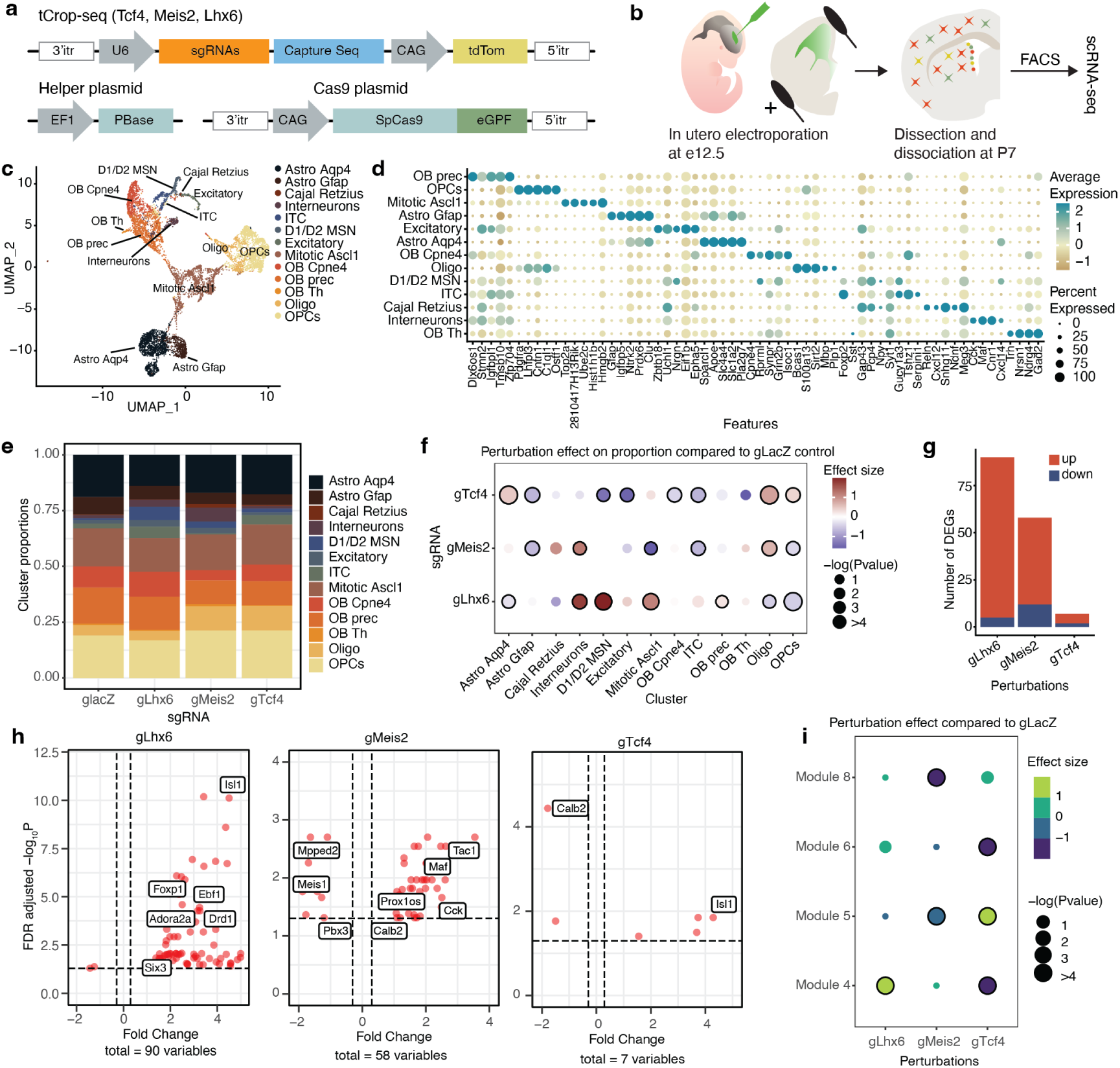
Embryonic disruption of developmental TFs alters postnatal cell types. Schematics of tCrop-seq vector maps **a** and the experimental workflow **b**. **c**, UMAP plot of the P7 data colored by cell type. **d**, Dot plot showing the top 5 marker genes of each cell type. OB, olfactory bulb cells; OPC, oligodendrocyte progenitor cells; ITC, intercalated cells; MSN, medium spiny neurons; Oligo, oligodendrocyte, Astro, astrocytes. **e**, Cell type compositions for each sgRNA. **f**, Perturbation effects in different clusters and sgRNAs compared to glacZ controls. Dot color corresponds to effect size, dot size corresponds to negative base 10 log(P-value). P-values were calculated from linear modeling, Padj was calculated by Benjamini & Hochberg FDR correction. The black outline indicates statistical significance (p-val < 0.05). **g**, Bar plot showing the number of differentially expressed genes detected in each sgRNA. **h**, Volcano plot showing differentially expressed genes in inhibitory neurons for each sgRNA, compared to gLacZ that meet the cut-off criteria (FDR < 0.05, avg_logFC > 0.5). **i**, Dot plot showing the effect of perturbation by sgRNAs on the module scores of inhibitory modules. The p-values were adjusted using Bonferroni correction.

ScRNA-seq data are highly heterogeneous and have numerous zero counts, making it challenging to detect subtler perturbation-based biological changes in single cell datasets. To overcome these limitations, we utilized Hotspot (DeTomaso and Yosef, 2021), a tool that identifies co-varying groups of genes (modules). Each cell was assigned a gene module score, with higher scores indicating higher association with that module. We identified 8 Hotspot gene modules (Figure S10a), 4 of which were neuronal (Figure 6i, S10b). Module 5 represented mostly in OB neuroblasts and contained genes enriched for neuronal differentiation. Module 4 represented MSN cell types and contained MSN marker genes (*e.g.*, *Foxp1*) and genes involved in retinoic acid receptor signalling (*Rarb*, *Rxrg*). The retinoic acid pathway is involved in the switch between proliferation and differentiation (Berenguer and Duester, 2022), which is essential for striatal development (Chatzi et al., 2011). Module 8 was represented in OB precursors and ITC cells. This module contained *Meis2*, as well as some of its target genes, such as *Pbx3* and *Etv1* (Table S4). Module 6 represented the OB-Cpne4 population and was characterized by genes involved in calcium response and synapse organization. We fitted a linear regression model that accounted for the batch and number of genes, and extracted the effect sizes to estimate how the module scores in the perturbed cells deviated from gLacZ control cells (Jin et al., 2020). For the three TFs, the perturbations had significant effects across different modules (FDR-corrected P < 0.05; Figure 6i). The perturbation of *Lhx6* was positively associated with the expression of module 4, consistent with the change in cell proportion and change in differentially expressed genes. The perturbation of *Meis2* lowered the expression of both modules 8 and 5. The perturbation of *Tcf4* had a significant effect across modules 6, 5, and 4, consistent with previous findings showing that TCF4 is a key facilitator of neurogenesis and neuronal differentiation (Figure 6i) (Mesman et al., 2020; Teixeira et al., 2021). Taken together, the tCROP-seq data at P7 indicate a marked influence of MEIS2, LHX6, and TCF4 on PN and IN specification.

## Discussion

### MEIS2 induces GABAergic projection neuron fate

In this study, we explored the role of the TF MEIS2 in the development of GABAergic PNs and INs in the mammalian telencephalon. Our study uses a new method combining transposon-based strategies for CRISPR perturbation sequencing (tCrop-Seq) and barcode lineage tracing (TrackerSeq). Consistent with a previous study, in which a conditional *Meis2* knockout mouse line was used (Su et al., 2022), CRISPR-induced perturbation of *Meis2* in the GE decreased the expression of PN-specific genes, and reduced the proportion of LGE-derived GABAergic PN types being generated. Along with this, we have observed an increase in the proportion of CGE-derived IN types being generated. We conducted *in vitro* reporter assays and found that MEIS2 requires the presence of DLX proteins to direct its functional activity towards regulatory enhancers of PN-specific genes containing specific co-binding sites.

### The spatial selective activation of enhancers

Our findings contribute to an overall picture in which spatial selective enhancer activation plays a role in the early imprinting of GABAergic identities (Figure S11). Different GABAergic cell types arise from regional differences in the specification of GE progenitors, which are initially established by morphogenic molecules such as retinoic acid (RA, LGE) (Chatzi et al., 2011), fibroblast growth factor (FGF) 8 and sonic hedgehog (SHH, MGE) (Storm et al., 2006; Molotkova et al., 2007), FGF12 and FGF15 (CGE) (Borello et al., 2008; Shohayeb et al., 2021), and their downstream TFs, such as MEIS2 (LGE), NKX2.1 & LHX6 (MGE), and NR2F1/2 (CGE). Our results depict how spatial factors are utilized downstream for selective enhancer activation: The tissue specificity of members of the DLX family in the GE, directs the functional activity of MEIS2 to regulatory sites related to GABAergic PN development. This is consistent with a proposed model of TALE TFs (e.g., MEIS) acting as broad co-activators of homeobox genes (Bridoux et al., 2020). Multiple studies have demonstrated that MEIS proteins require the presence of other TFs, such as PBX, HOX, TBX, and PAX6, to promote differentiation in the limbs, heart, lens, hindbrain, and olfactory bulb (Schulte and Geerts, 2019; Bridoux et al., 2020; Delgado et al., 2021; Selleri et al., 2019; Agoston et al., 2014). Furthermore, MEIS2 appears to act in a highly context-dependent manner, as evidenced by the minimal overlap between MEIS1/2 ChIP-seq data from the retina (Dupacova et al., 2021) and ChIP-seq data from the GE (data not shown).

DLX/MEIS2 could inhibit IN fate through the activation of repressive transcription factors, such as ISL1, FOXP1/2, and SIX3, via co-repressors such as TLE1/4, or by promoting the expression of microRNAs (miRNAs). We identified several miRNA host genes that were downregulated in Meis2-tCROP-seq: *Mir124-2hg Gm27032*(miR-124a-2), *Arpp21* (miR-128-2), and *Gm27032* (miR-124a-3; Table S4). miR-124 and miR-128 are some of the most abundant and highest enriched miRNAs in the adult mouse and human brains (Zolboot et al., 2021). miR-128 deficiency in D1-MSNs leads to juvenile hyperactivity, followed by lethal seizures at 5 months of age (Tan et al., 2013).

### LHX6 plays an antagonistic role to MEIS2

We demonstrated that in GE, MEIS2 and DLX5 together activate several enhancers associated with PN gene expression that we tested and that are active in the LGE (Figure S7). This spatial component appears to be partially mediated by LHX6, which antagonizes MEIS2 in two ways: First, we show that LHX6 suppresses a regulatory element of *Meis2*, likely resulting in repression of *Meis2* gene expression in the SVZ/MZ of the MGE. Consistently, *Meis2*, as well as the PN marker genes *Pbx3* and *Foxp1*, have been shown to be up-regulated in E14.5 *Lhx6* knockout cells collected from the cortex (Asgarian et al., 2022). Furthermore, conditional knockout of *Nkx2-1*, which acts upstream of LHX6, has been shown to result in increased transcription of *Meis2* in the SVZ of the MGE (Sandberg et al., 2016) and an enrichment of repressive regulatory elements in motifs consistent with the binding site of MEIS2 (Sandberg et al., 2018). Second, while *Meis2* mRNA is rapidly downregulated as cells enter the SVZ/MZ, MEIS2 protein decay is expected to be slower (Figure S6) (Fischer et al., 2014). We found co-immunoreactivity of MEIS2 and LHX6 in a small population of cells around the interface of the VZ and SVZ in the MGE (Figure 5a). To counteract the residual protein activity, we found that LHX6 can efficiently repress the cooperative MEIS2/DLX5 activation of PN fate genes in the MGE (Figure 5f,g,i, S3e). The suppression by LHX6 could be mediated by a competition of LHX6 with DLX for the common DNA binding motif TAATT (Sandberg et al., 2016; Lindtner et al., 2019). Alternatively, LHX6 could restrict the interaction of MEIS2/DLX5 with DNA through direct binding to DLX5 or MEIS2. LHX6 belongs to the LIM domain homeodomain (LIM-HD) protein family, which is characterized by two cysteine-rich LIM domains for protein-protein interactions and a homeodomain for binding DNA (Hobert and Westphal, 2000). For example, LHX6 directly interacts with PITX2 to inhibit its transcriptional activities (Zhang et al., 2013). In parallel, other transcriptional programs are likely involved in the repression and activation of PN and IN cell fate (Chapman et al., 2018; Leung et al., 2022).

### Differential binding or differential accessibility?

Delas et al. 2023 (Delás et al., 2023) propose two *cis*-regulatory strategies that could drive cell fate choice in developing neural progenitors: One - differential binding - relies on a common regulatory landscape, whereby the different composition of TFs at these *cis*-regulatory elements dictates differential gene expression and cell fate decisions. The other - differential accessibility - relies on cell-type-specific chromatin remodeling. Our results support the first strategy: The enhancers we studied, are accessible in both the LGE and the MGE, regardless of their *in vivo* activity pattern (Figure S7). Furthermore, while the selected enhancers are accessible throughout all GEs, our data show that their activity depends on the TF composition. For example, the *Foxp2* enhancer “hs956” is not active in the VZ of the GE (Figure S7), likely because *Dlx* genes are absent in the VZ. This enhancer is active in the SVZ and MZ of the LGE, where both *Meis2* and *Dlx* genes are expressed. The enhancer is not active in the SVZ/MZ of the MGE where *Meis2* is absent, and a repressive TF such as *Lhx6* is present (Figure S6, S7).

### How do MEIS2 and DLX5 work together?

Agoston et al. 2014 (Agoston et al., 2014) performed pull-down experiments with a tagged form of MEIS2 using OB tissue, and detected DLX-specific protein bands in the MEIS2 precipitates. This could either indicate a direct protein interaction or be the result of a process called ’DNA-guided cooperativity’, a mechanism where certain TFs cooperatively bind to adjacent DNA sites without forming stable, direct protein-protein interactions (Kim et al., 2023). This form of co-binding is guided by the DNA sequence itself, rather than by protein-protein interactions. In support of “DNA-guided cooperativity” as the mechanism underlying the interaction between MEIS1 and DLX3, is a study by Jolma et al. (Jolma et al., 2015), which performed *in vitro* structural analysis of the TF pairs, included a crystal structure of MEIS1 and DLX3 bound to their identified recognition site. Their results demonstrated that the interactions between MEIS and DLX are predominantly mediated by DNA.

### MEIS2 in Pathology

Haploinsufficiency of the MEIS2 in humans results in an autosomal dominant disease characterized by multiple congenital malformations, mild-to-severe intellectual disability with poor speech, and delayed psychomotor development (Louw et al., 2015; Douglas et al., 2018; Giliberti et al., 2020; Gangfuß et al., 2021; Zhang et al., 2021). The amino acid Arg333, located in the homeodomain of MEIS2, is highly conserved across species and isoforms (Longobardi et al., 2014), and was found mutated in at least four patients with severe disease (Giliberti et al., 2020; Gangfuß et al., 2021). Our study found that the missense mutation p.Arg333Lys led to a strong decrease in enhancer activation compared to wild-type MEIS2. Due to the location of Arg333 in the homeodomain of MEIS2, it is likely that the mutations in this amino acid interfere with the protein’s DNA binding ability. This could result in a change in GABAergic cell type proportions, in particular a reduced number of PNs in the striatum, caused by disturbed fate decisions during embryogenesis, and ultimately elicit the disease phenotype seen in affected individuals.

## Conclusion

The efficiency with which MEIS2 can co-activate selective enhancers suggests a general strategy for implementing spatial information to generate distinct cellular populations. The ability of MEIS2 to induce context-specific cell types may exemplify how certain subsets of cells in different parts of the body are affected in developmental disorders. Further research is needed to fully comprehend the intricate interactions between TFs and co-factors in the regulation of cell fate decisions during GABAergic neuron development and their potential implications in human disease.

## Data availability

The datasets used in this research article can be downloaded from the Gene Expression Omnibus (GEO) accession number GSE231779 (secure reviewer access token: cpclgssspnqpvmv). Additionally, the code to reproduce the data analysis is available at .

## Acknowledgements

We thank members of the Mayer and Winkelmann laboratories for feedback and discussion; J. Kuhl (somedonkey.com) for illustrations; R. H. Kim from the MPIB Next-Generation Sequencing core facility, I. Velasques and G. Eckstein from the Genomics Core facility at the Helmholtz Zentrum Munich (HMGU), M. Spitaler and M. Oster from the MPIB Imaging and FACS core facility, R. Kasper from the MPIBI Imaging core facility and members of the MPIB/MPIBI animal facility for their technical expertise. This work was supported by the Max-Planck Society, the European Research Council (ERC) under the European Union’s Horizon 2020 Research and Innovation program (ERC-2018-STG, grant agreement no. 803984, GIDE; to C.M.), and the European Commission (SMART GRANT: ERA-NET NEURON, SMART: 01EW1605).

## Author contributions statement

E.D. and C.M. conceived the project; M.H. and C.M. developed TrackerSeq; E.D. and C.M. developed tCROP-seq; E.D., I.V. and C.M. conducted the tCROP-seq and TrackerSeq experiments; D.D.L., I.D. and M.T. conducted the MEIS1/2 ChIP-seq experiments; E.D. conducted functional reporter assay experiments; M.H. lead scRNA-seq, tCROP-seq, TrackerSeq computational analyses; V.K. lead ChIP-seq analyses; F.N. conducted the logistic regression analysis; E.D, M.H., V.K., F.N., J.W. and C.M. prepared the manuscript with input from the remaining authors. **Competing interests:** The authors declare no competing interests.

## Supplementary Figures

**Supplementary Figure 1:**
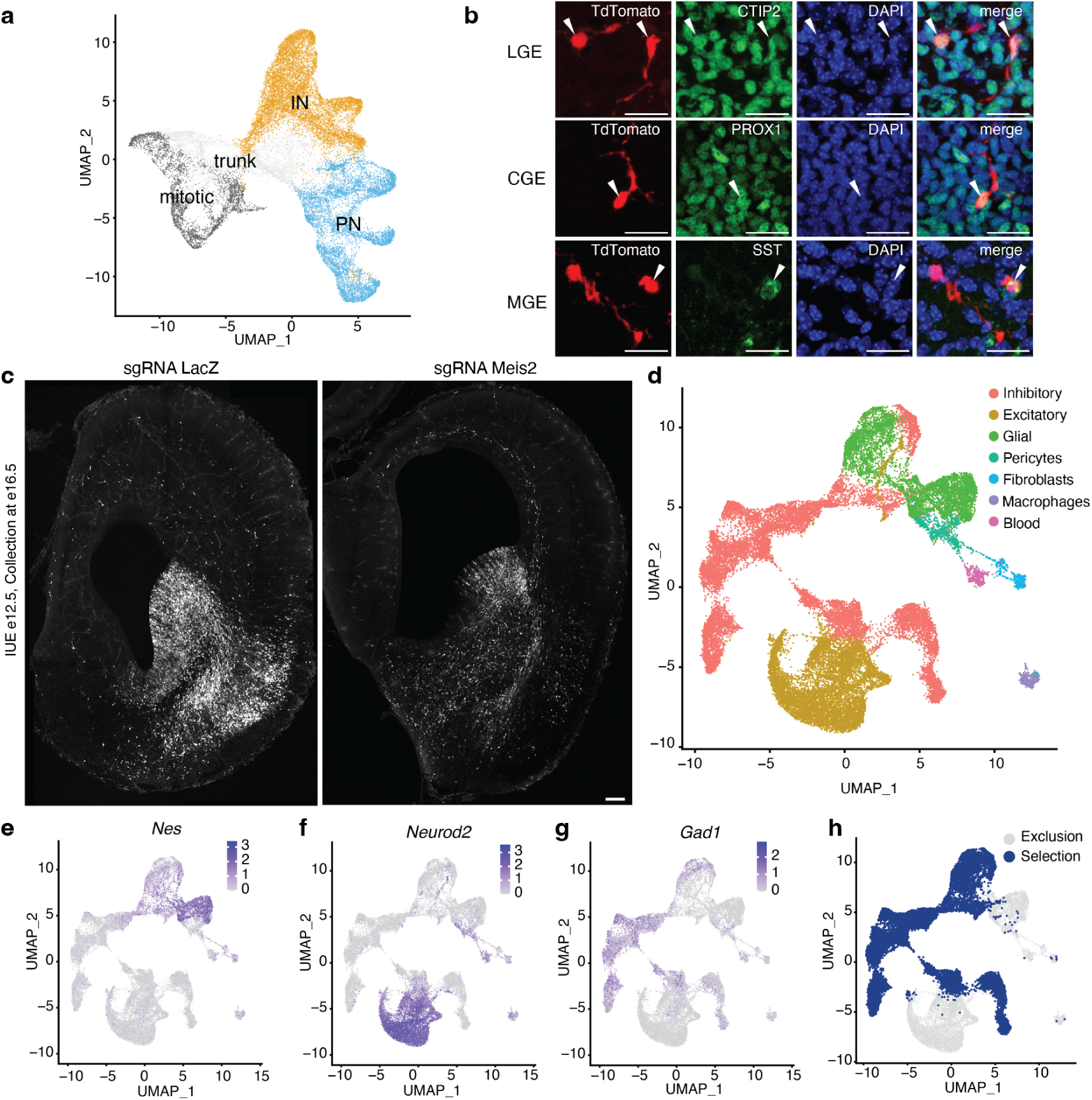
*In utero* electroporation of sgRNA targets subtypes of GABAergic neurons. **a**, UMAP depicting groups of cells used for the logistic regression analysis to predict PN and IN fate genes. Date from Bandler et al., 2022 (Bandler et al., 2022). **b**, Immunohistochemistry of E18.5 brains electroporated with tCROP-seq LacZ sgRNA vector at E12.5. Subsets of tdTomato-expressing neurons show immunoreactivity against markers of different inhibitory neuron types: anti-CTIP2, LGE-derived striatal PNs; anti-SST, MGE-derived cortical INs; anti-PROX1, CGE-derived cortical INs. **c**, Localization of tdTomato expression driven by gLacZ and gMeis2 plasmids in the cortex, striatum, and GE at E16.5, following IUE at E12.5. Scale bar, 0.1 mm. **d**, UMAP plot displaying E16 data colored according to cell classes. **e-g**, Feature plots depicting the expression of the canonical marker genes *Nes*, *Neurod2*, and *Gad1*. **h**, UMAP plot illustrating the selection of cells for downstream analysis.

**Supplementary Figure 2:**
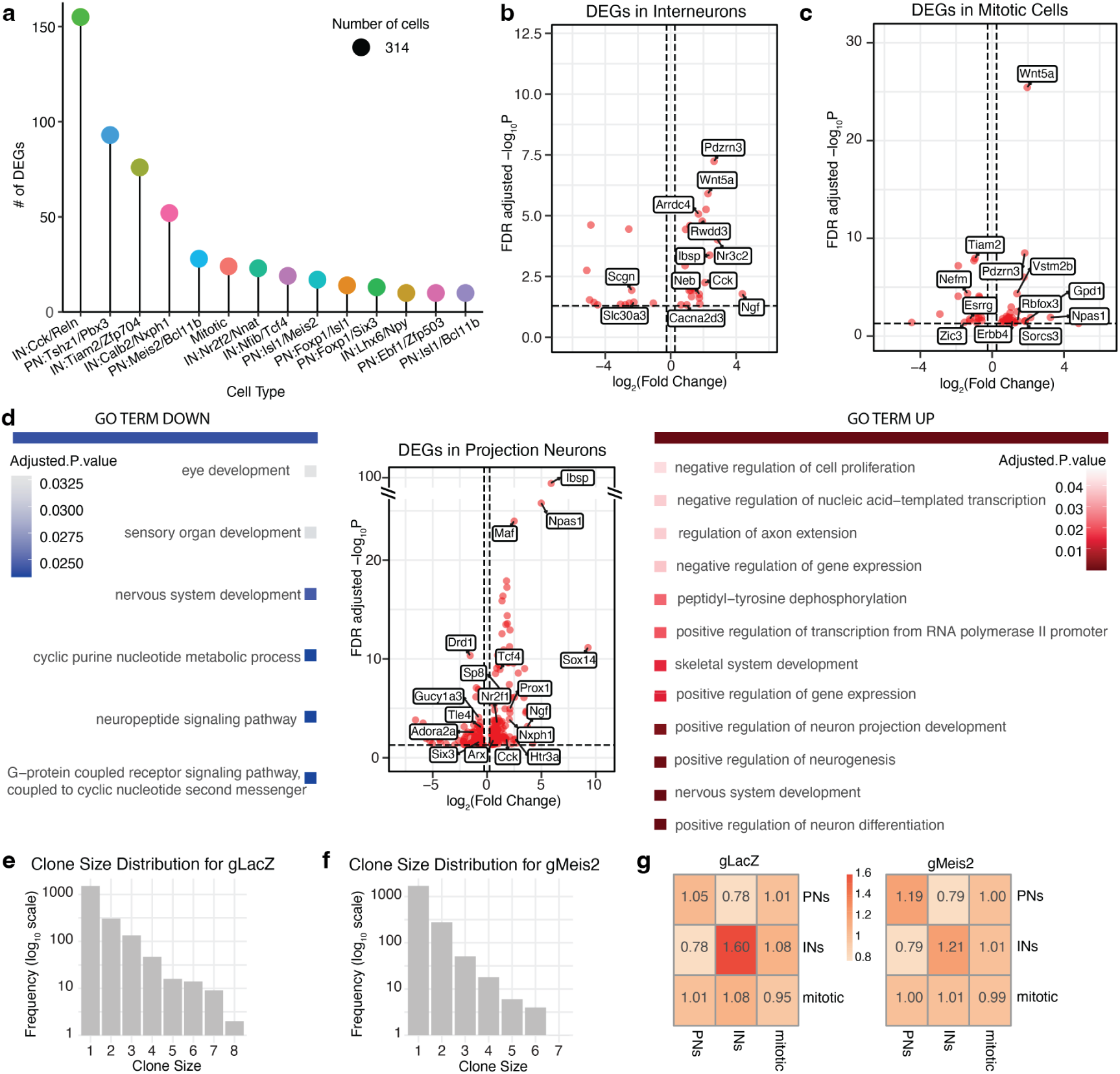
gMeis2 modulates gene expression and influences clonal coupling. **a**, Lollipop plots illustrating the impact of gMeis2 on inhibitory clusters, with the number of differentially expressed genes (DEGs) shown after downsampling each group to 314 cells. **b**, Volcano plot depicting the differentially expressed genes in gMeis2 and gLacZ interneurons. **c**, Volcano plot depicting the differentially expressed genes in gMeis2 and gLacZ mitotic cells. **d**, Gene ontology analysis on differentially expressed genes (DEG) of clusters belonging to the projection neuron class. **e**, Histogram illustrating the distribution of clone sizes for the gLacZ TrackerSeq dataset. **f**, Histogram illustrating the distribution of clone sizes for the gMeis2 TrackerSeq dataset. **g**, Clonal overlap between cell states. The number of shared barcodes between pairs is normalized by expectation if clonal membership is shuffled.

**Supplementary Figure 3:**
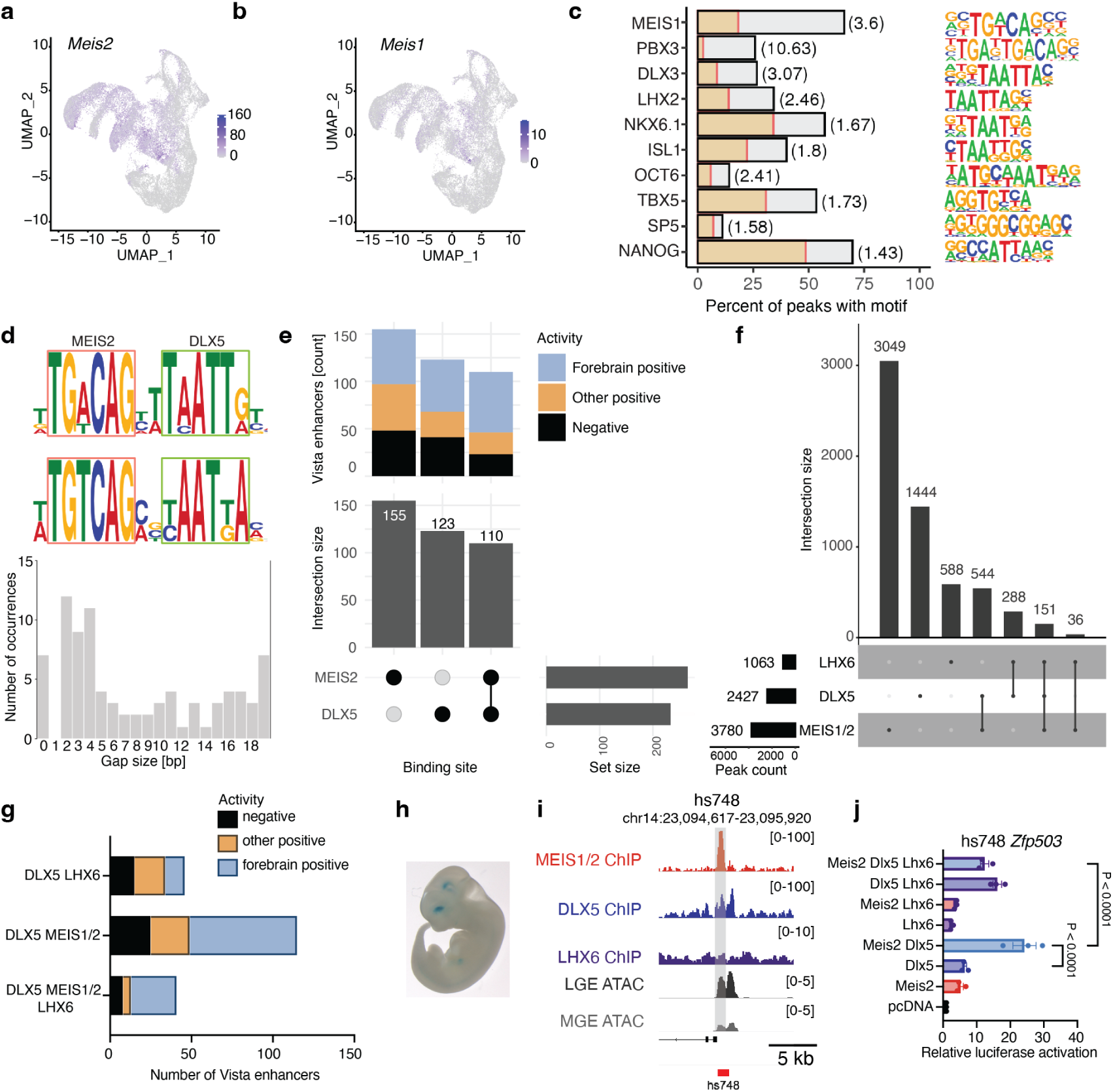
Overlap of Meis1/2 and Dlx5 ChIP-seq binding sites in the ganglionic eminence (GE). **a**, Feature plot depicting the expression level of *Meis2* at E16. **b**, Feature plot depicting the expression level of *Meis1* at E16. **c**, Motif occurrence analysis of selected known motifs enriched within all MEIS1/2 binding sites (grey bars) compared to G/C-matched reference sequences (yellow). **d**, Motif spacing analysis of MEIS2 and DLX5 motifs within shared binding sites. The position weight matrix (PWM) of the most frequent motif configuration is shown on the left, while the right panel illustrates the overall distribution of the DLX5 motif in relation to the MEIS2 motif. **e**, Overlap analysis of binding sites between MEIS1/2 and DLX5 (bottom) and their distribution within different classes of Vista enhancers (top). **f**, Overlap analysis of binding sites between MEIS1/2, DLX5, and LHX6. **g**, Visualization of LacZ expression driven by the hs748 enhancer in the E12.5 mouse forebrain (Visel et al., 2007). **h**, Representative tracks of GE ChIP-seq of MEIS1/2 at E14.5 (red), DLX5 at E13.5 (blue) (Lindtner et al., 2019), LHX6 at E13.5 (purple) (Sandberg et al., 2016) and scATAC-seq (Rhodes et al., 2022) from the LGE (dark gray) and MGE (gray) at E12.5. **i**, Overlap between binding sites of MEIS1/2, DLX5, and LHX6 in enhancer hs748, which associated with the gene *Zfp503*. **j**, Luciferase activity driven by hs748, co-transfected with *Meis2*, *Dlx5*, and *Lhx6* expression vectors in Neuro2a cells. Bars represent mean ± s.e.m from a total of 9 replicates, split into three independent batches, each performed in triplicate. Points represent the mean of each batch for each condition. Statistical significance was assessed by two-way ANOVA. P-values of pairwise comparisons from post-hoc Tukey’s HSD are presented for selected conditions.

**Supplementary Figure 4:**
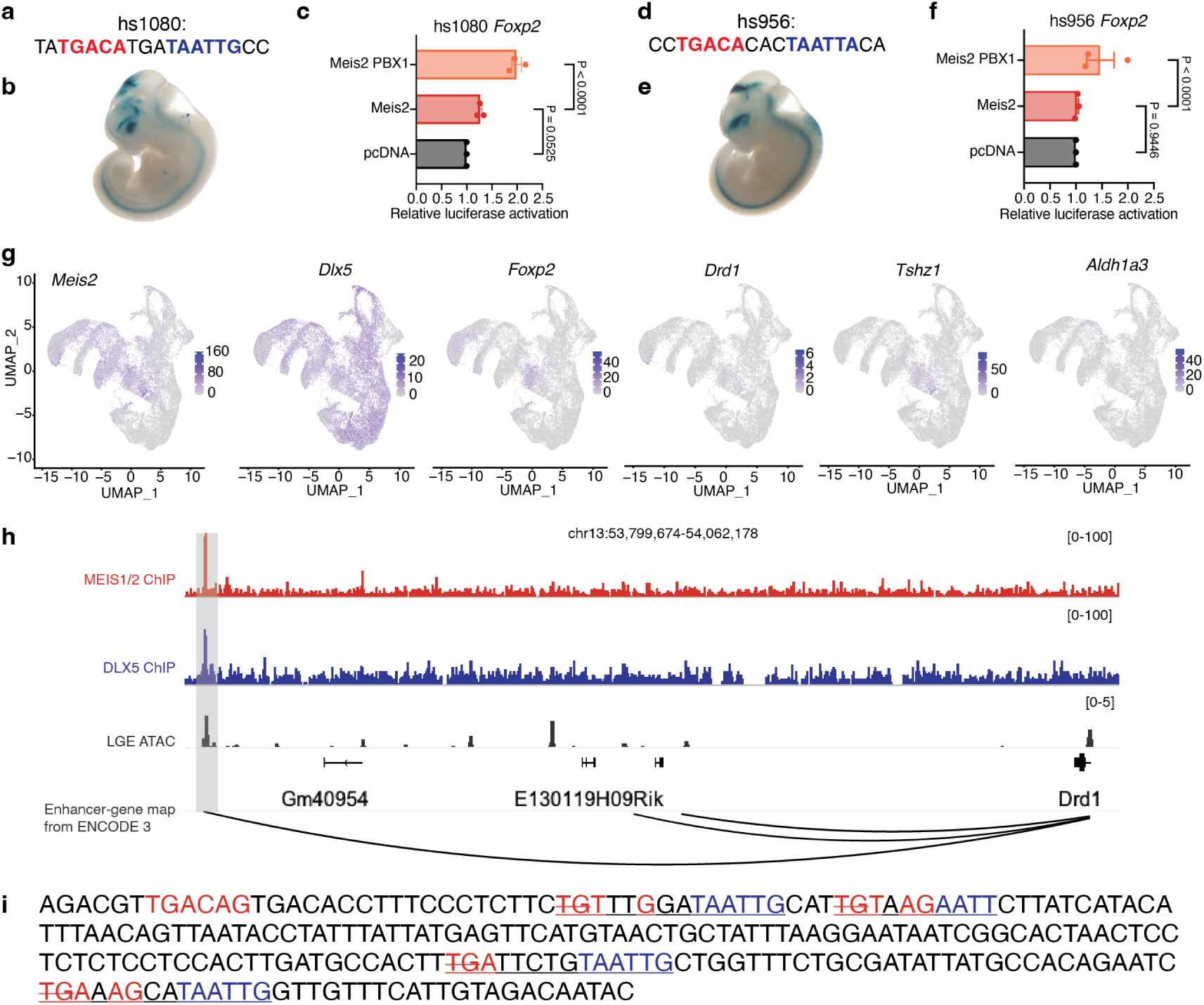
Regulation and functional analysis of PN enhancers. **a**, **d**, Combined MEIS (red) and DLX (blue) binding motifs found within hs1080 (a) and hs956 (d) enhancers. **b**, hs1080 and **e** hs956 enhancers drive LacZ expression in E12.5 mouse forebrain (Visel et al., 2007). **c**, **f** Luciferase assay measuring the activation effect of MEIS2 and PBX1 on hs1080 (c) or hs956 (f) driven luciferase reporter in Neuro2a cells. **g**, Feature plot depicting the expression level of *Meis2*, *Dlx5*, *Foxp2*, *Drd1*, *Tshz1* and *Aldh1a3* at E16. **h**, Visualization of the *Drd1* locus with aligned tracks of MEIS1/2 ChIP-seq at E14.5 (red), DLX5 ChIP-seq at E13.5 (blue), and LGE (dark grey) (Rhodes et al., 2022). The predicted enhancer-gene interactions are also depicted (Gorkin et al., 2020). **i**, Depiction of the DNA sequence of the shortened version of the enhancer enhD1, highlighting the combined MEIS (red) and DLX (blue) binding motifs. The TG bases removed in the mutated version of enhD1 are indicated with a strikeout line. In panels c and f, bars represent mean ± s.e.m from a total of 9 replicates, split into three independent batches, each performed in triplicate. Points represent the mean of each batch for each condition. Statistical significance was assessed by two-way ANOVA. P-values of pairwise comparisons from post-hoc Tukey’s HSD are presented for selected conditions.

**Supplementary Figure 5:**
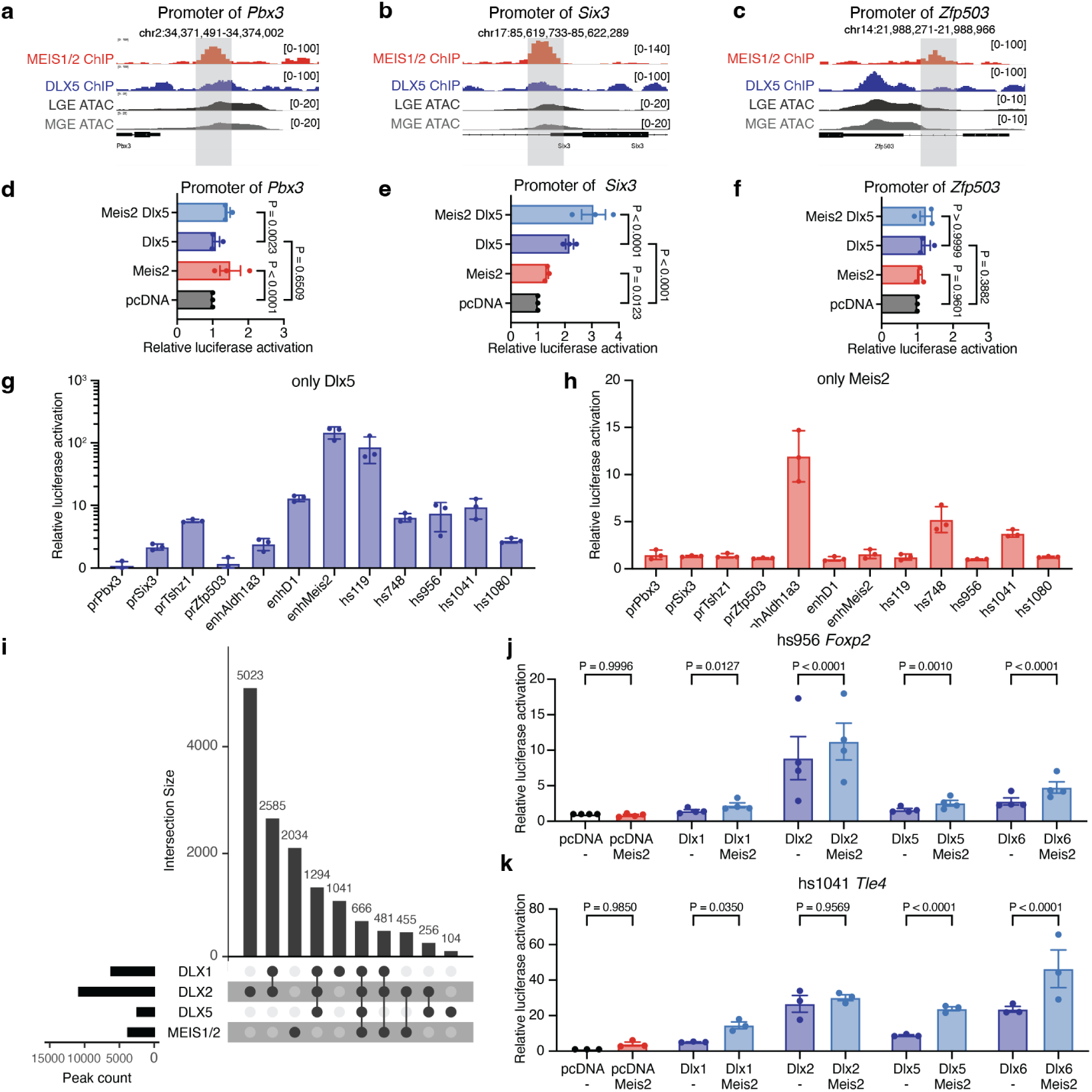
MEIS2 acts primarily via distal enhancers in the GE. **a-c**, Representative tracks of MEIS1/2 ChIP-seq in the GE at E14.5 (red), DLX5 ChIP-seq in the GE at E13.5 (blue) (Lindtner et al., 2019), and scATAC-seq in the LGE (dark gray) and MGE (gray) at E12.5 (Rhodes et al., 2022) are shown at the gene promotors of *Pbx3*, *Six3* and *Zfp503*. **d-f**, Luciferase activity driven by promoters of *Pbx3*, *Six3* and *Zfp503* genes, transfected with MEIS2 and DLX5 expression vectors in Neuro2a cells. **g-h**, Luciferase activity driven by regulatory elements, transfected with MEIS2 (**g**) or DLX5 (**h**) expression vectors in Neuro2a cells. The data represents the combined results from multiple experiments. **i**, Overlap between binding sites of MEIS1/2, DLX1, DLX2 and DLX5. **j-k**, Luciferase activity driven by regulatory elements hs956 (**j**) and hs1080 (**k**), transfected with MEIS2 (**g**) and DLX1, DLX2, DLX5 or DLX6 expression vectors in Neuro2a cells. In panels d, e, f, g, h, j and k, bars represent mean ± s.e.m from a total of 9 or 12 replicates, split into 3-4 independent batches, each performed in triplicate. Points represent the mean of each batch for each condition. Statistical significance was assessed by two-way ANOVA. P-values of pairwise comparisons from post-hoc Tukey’s HSD are presented for selected conditions.

**Supplementary Figure 6:**
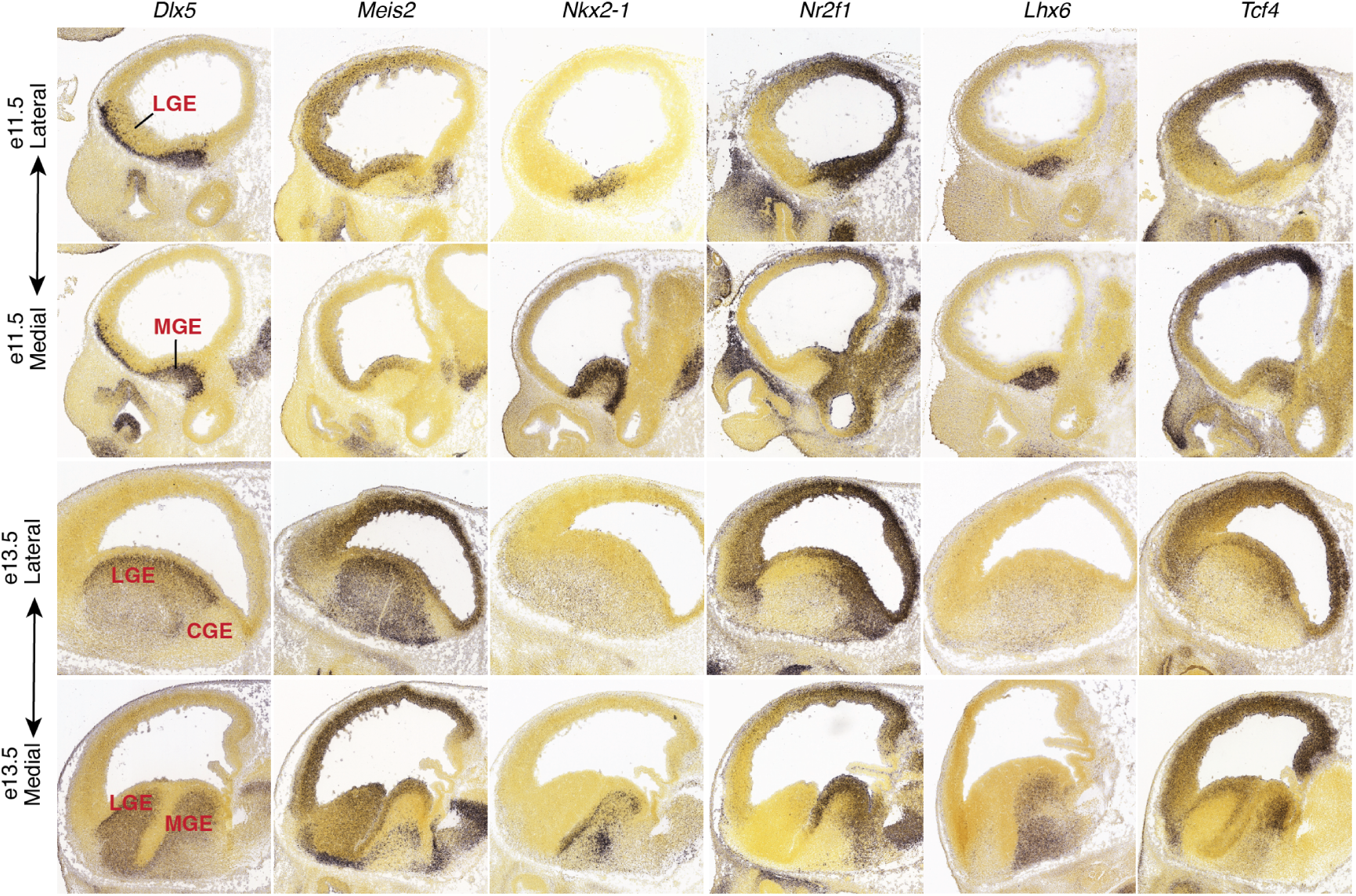
Spatial expression patterns of TFs in the GE. *In situ* Hybridization (ISH) images of *Dlx5*, *Meis2*, *Nkx2-1*, *Nr2f1*, *Lhx6*, and *Tcf4* from the Allen Brain Institute’s Developing Mouse Brain Atlas at E11.5 and E13.5. MGE, medial ganglionic eminence; LGE, lateral ganglionic eminence; CGE, caudal ganglionic eminence) are indicated.

**Supplementary Figure 7:**
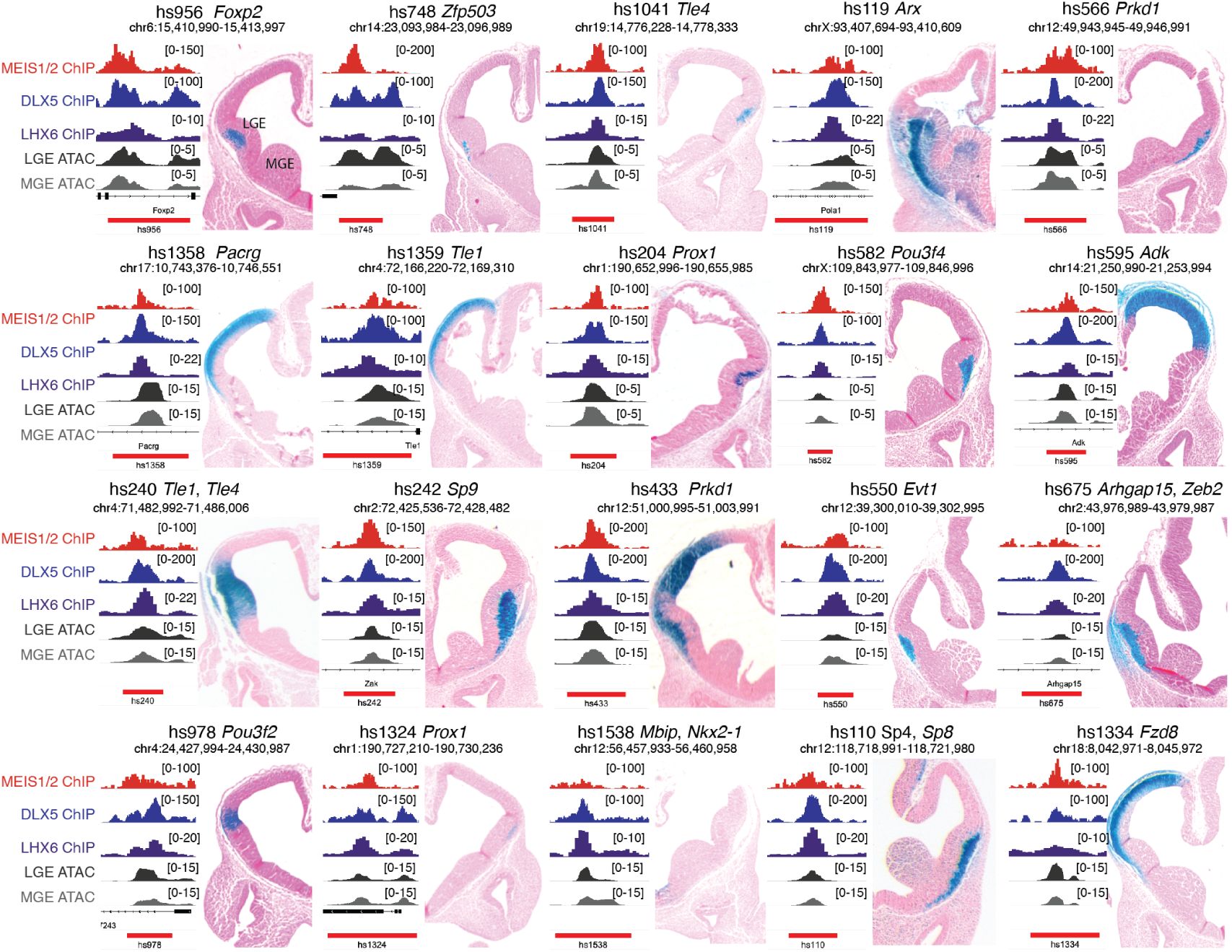
Spatial activity of Meis2 targets. Selected Vista enhancers with *in vivo* activity at E11.5 (Visel et al., 2007) and co-binding of MEIS-DLX5-LHX6. On the left side of each image are panels with representative tracks of GE ChIP-seq of MEIS1/2 at E14.5 (red), DLX5 at E13.5 (blue) (Lindtner et al., 2019), LHX6 at E13.5 (purple) (Sandberg et al., 2016) and scATAC-seq (Rhodes et al., 2022) from the LGE (dark gray) and MGE (gray) at E12.5. MGE, medial ganglionic eminence; LGE, lateral ganglionic eminence.

**Supplementary Figure 8:**
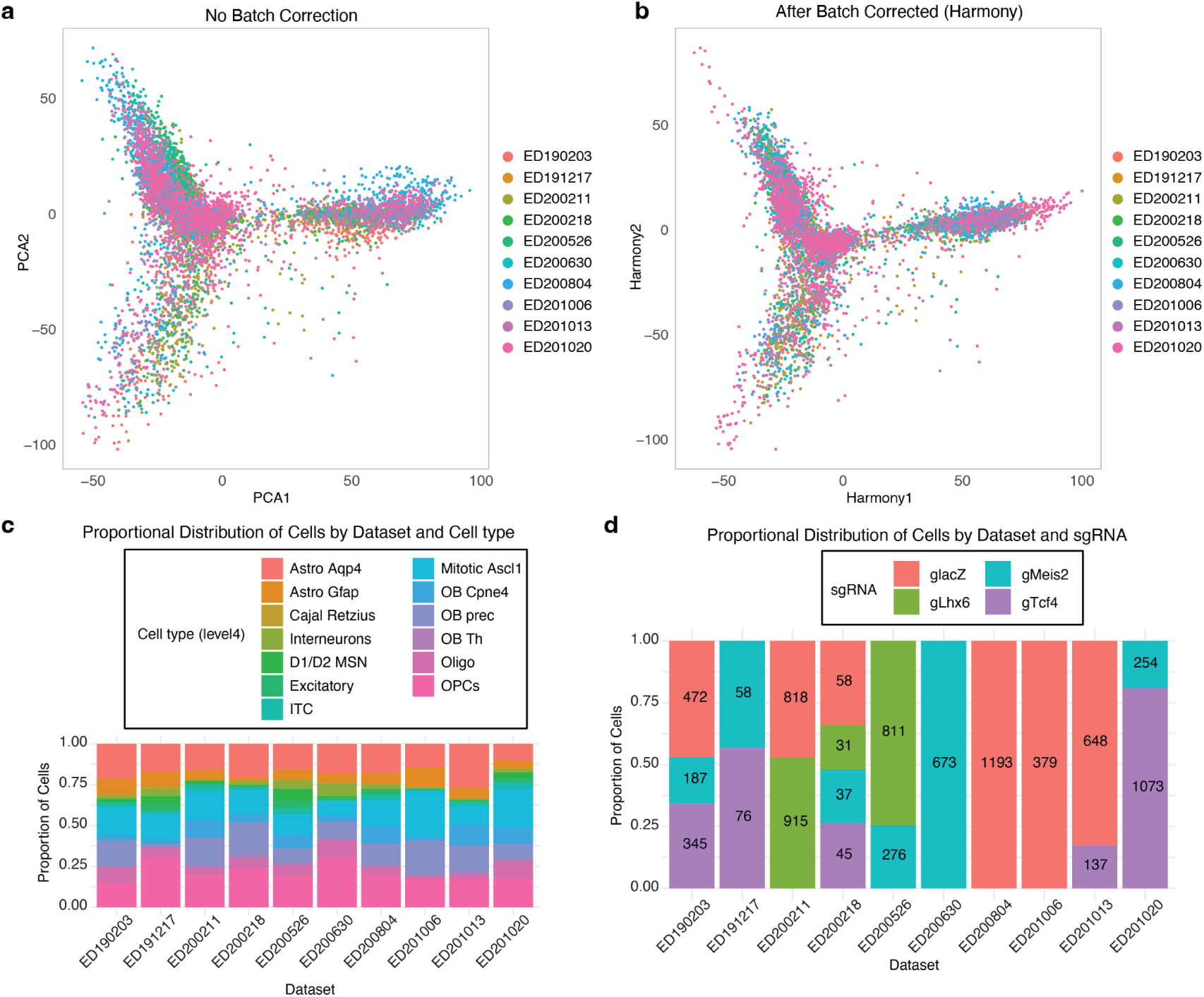
Batch correction and sgRNA coverage of P7 tCROP-seq datasets. **a-b**, 2D visualization of the P7 tCROP-seq dataset pre (a) and post batch correction using Harmony (Korsunsky et al., 2019). **c**, Proportional distribution of cells categorized by dataset and cell type for the P7 tCROP-seq dataset. **d**, Proportional distribution of cells categorized by dataset and sgRNA for the P7 tCROP-seq dataset.

**Supplementary Figure 9:**
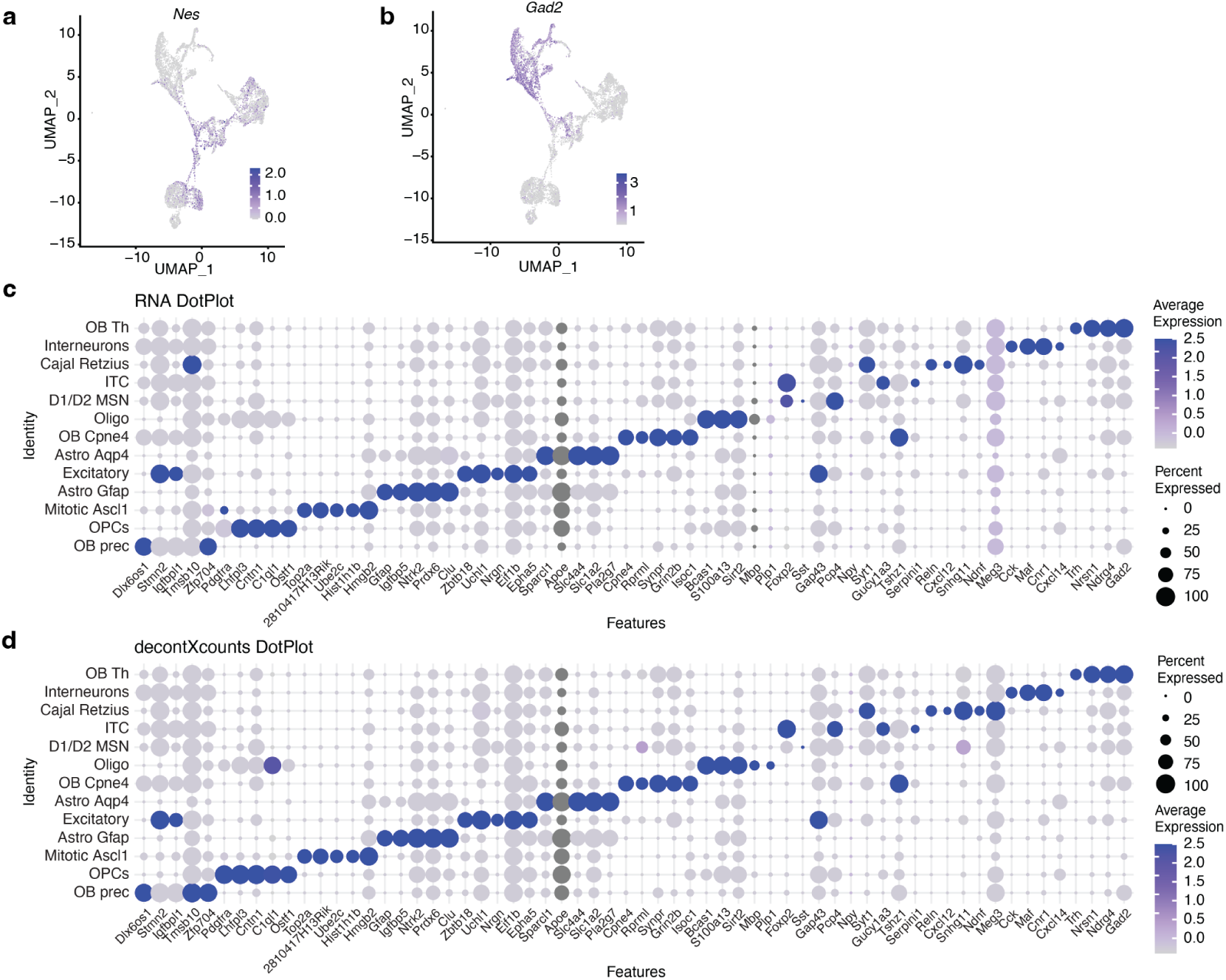
Correct of the P7 tCROP-seq data for non-specific background expression. **a-b**, Feature plots of canonical marker genes *Gad2* and *Nes* at P7. **c**, Dotplot demonstrating the top marker genes of inhibitory clusters using the “RNA count” data. **d**, Dotplot illustrating the top marker genes of inhibitory clusters using the “decontXcounts” (Yang et al., 2020) data, which corrects for potential non-specific background expression. The differences before and after correction are small.

**Supplementary Figure 10:**
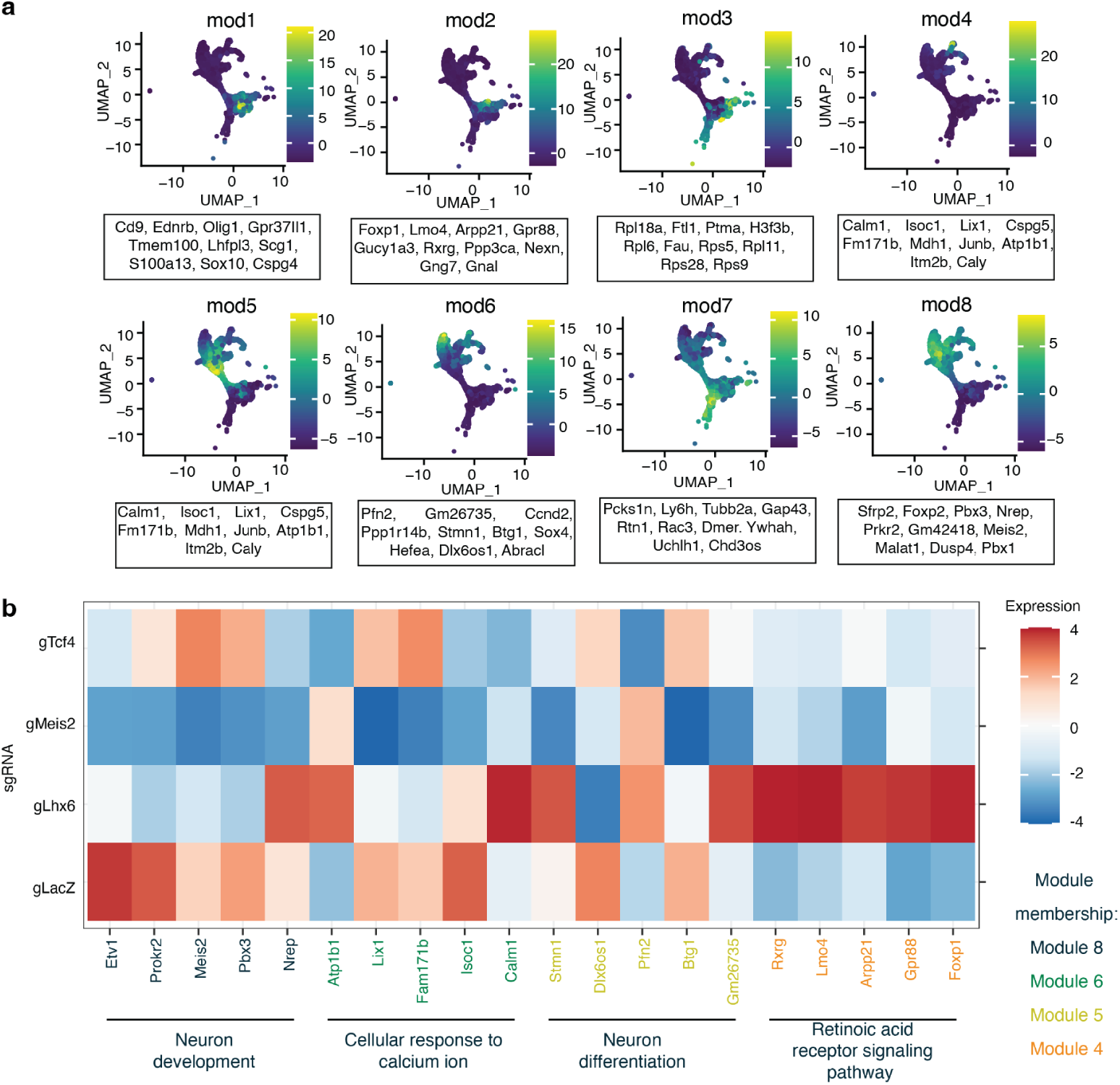
Module analysis using the P7 tCROP-seq dataset. **a**, Feature plots of gene module expression scores and the correlated genes within each module. **b**, Average expression of top 5 module genes for each sgRNA at P7.

**Supplementary Figure 11:**
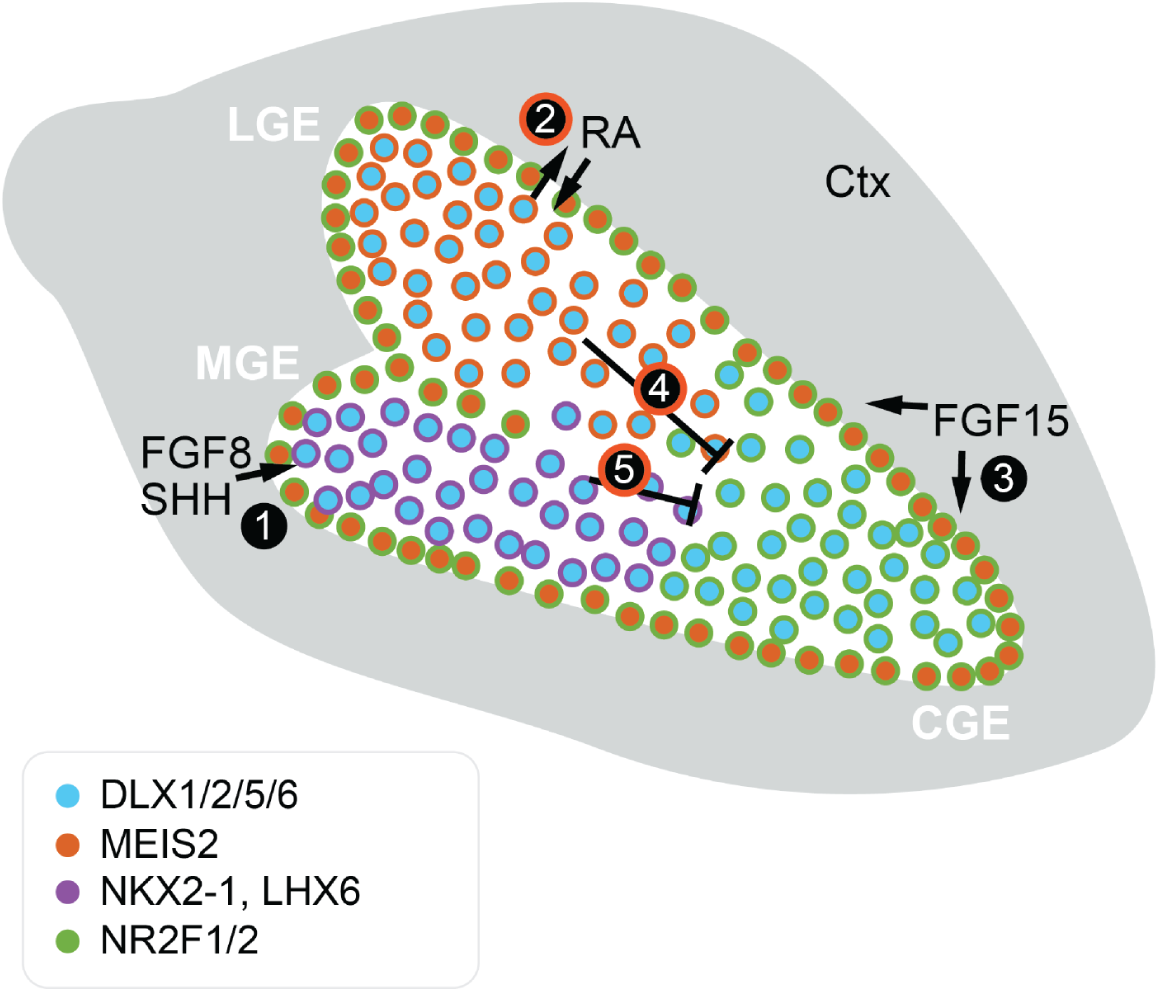
Schematic summary of spatial factors in the ganglionic eminence leading to specific enhancer activation. MGE, medial ganglionic eminence; CGE, caudal ganglionic eminence; LGE, lateral ganglionic eminence; Ctx, cortex; RA, retinoic acid; SHH, sonic hedgehog; FGF, fibroblast growth factor. 1. (Storm et al., 2006; Molotkova et al., 2007); 2, (Chatzi et al., 2011); 3, (Borello et al., 2008; Shohayeb et al., 2021); 4, (Su et al., 2022); 5, (Vogt et al., 2014). The red circle outline represents the findings of this study.

## Supplemental Material

Tables are presented as individual Excel files.

- Table S1: Selected sgRNAs list with primers plus TrackerSeq primers.
- Table S2: E16-tCrop-seq top10 marker genes.
- Table S3: E16-tCrop-seq pseudo-bulk differential gene expression analysis.
- Table S4: E16-tCrop-seq interneuron clusters differential gene expression analysis.
- Table S5: ChIP-seq supplementary information.
- Table S6: P7-tCrop-seq top10 marker genes.
- Table S7: P7-tCrop-seq differential gene expression analysis.
- Table S8: Information on cloned regulatory elements used in luciferase reporter assays and detailed statistics.
- Table S9: Datasets information.

## Methods

### Cell line

Mouse Neuro2a neuroblastoma cells (ECACC, 89121404) were cultured in Dulbecco’s modified Eagle medium (DMEM, Sigma, D6429) with high glucose, L-glutamine, and sodium pyruvate supplemented with 10% (v/v) fetal bovine serum (FBS, Sigma, F9665) and containing 1% (v/v) antibiotics (100 U/mL penicillin, 100 mg/mL streptomycin) (Sigma, P0781). Neuro2a cells were incubated at 37 ℃in a 5% CO_2_ humidified atmosphere and passaged twice a week. Cell passage numbers were limited to no more than 10.

### sgRNA selection and vector construction

The piggyBac based backbone plasmid contains sgRNAs under the mouse U6 promoter, a gift from Randy Platt, were modified by adding pCAG-TdTomato (Addgene, 59462) and a capture sequence at the scaffold of sgRNA for 10x feature barcode retrieval (cs1 incorporated at the 3’ end; (Replogle et al., 2020)) with use of NEBuilder HiFi DNA Assembly (NEB, E2621). sgRNAs were designed using CRISPick for CRISPRko (Doench et al., 2016; Sanson et al., 2018) and validated with inDelphi (Shen et al., 2018) for high frame shift efficiency. At least 3 sgRNAs per gene were cloned using ssDNAs oligoes (IDT) and NEBuilder HiFi DNA Assembly (NEB) into modified backbone. The efficiency of sgRNA was measured in Neuro2A cells. Cells were transfected with pCAG-Cas9-EGFP (gift from Randy Platt) and sgRNAs plasmids with TransIT-LT1 Transfection Reagent (Mirus, MIR2305) and after 48 h were sorted with BD FACSAria III Cell Sorter (BD FACSDiva Software, version 8.0.2) for TdTomato and EGFP. The genomic DNA was extracted with Quick-DNA Miniprep Plus Kit (Zymo, D4068) and the region around sgRNA targeting was amplified with Q5 polymerase (NEB, M094S) with primers listed in the Table S1, and afterwards sent for Sanger sequencing at Microsynth Seqlab GmbH. The knockout efficiency quantified using the Synthego ICE Analysis Tool (Hsiau et al., 2019). The results for selected sgRNAs are shown in the Table S1.

### TrackerSeq library preparation and validation

TrackerSeq is a piggyBac transposon-based (Ding et al., 2005) library, which was previously developed by (Bandler et al., 2022) to be compatible with the 10x single-cell transcriptomic platform. It records the *in vivo* lineage history of single cells through the integration of multiple oligonucleotide sequences into the mouse genome. Each of these individual lineage barcodes is a 37-bp long synthetic nucleotide that consists of short random nucleotides bridged by fixed nucleotides. This design results in a library with a theoretical complexity of approximately 4.3 million lineage barcodes (16^8^) with each barcode differing from another by at least 5 bp. To construct the library, the piggyBac donor plasmid (Addgene, 40973) was altered to include a number of modifications (Bandler et al., 2022). A Read2 partial primer sequence was cloned into the 3’ UTR of the EGFP to enable retrieval by the 10x platform. The sucrose gene was cloned into the vector, so that empty plasmids that fail to incorporate a lineage barcode during the cloning process are removed. Following digestion with BstXI (Jena Bioscience, EN-E2118) to remove the sucrose gene, the plasmid was run on a gel and column purified. The lineage barcode oligo mix was cloned downstream of the Read2 partial primer sequence in the purified donor plasmid via multiple Gibson Assembly reactions (NEB, E2611S). Gibson assembly reactions were then pooled and desalted with 0.025 µm MCE membrane (Millipore, VSWP02500) for 40 min, and finally concentrated using a SpeedVac. 3 µl of the purified assembly is incubated with 50 µl of NEB 10-*β*-competent E.coli cells (NEB, C3019H) for 30 min at 4 ℃, then electroporated at 2.0 kV, 200 Ω, 25 µF (Bio-Rad, Gene Pulser Xcell Electroporation Systems). Electroporated E.coli were incubated for 90 min shaking at 37 ℃and then plated into pre-warmed sucrose/ampicillin plates. The colonies were scraped off the plates after 8 h, and the plasmids were grown in LB medium with ampicillin up to OD = 0.5. The plasmid library was purified using a column purification kit (Zymo, D4202). We first assessed the integrity of the TrackerSeq barcode libraries by sequencing the library to a depth of approximately 42 million reads to test whether any barcode was over-represented. Around 3.6 million valid lineage barcodes that had a quality score of 30 or higher were extracted from the R2 FASTQ files using Bartender (Zhao et al., 2018). One thousand barcodes were randomly sampled from the extracted lineage barcodes to assess hamming distance. To group similar extracted barcodes into putative barcodes, Bartender assigns a UMI to each barcode read to handle PCR jackpotting errors, and clusters them. The cluster distance was set to 3 so that extracted barcodes within 3 bp of each other have a chance of being clustered together. A total of 2 × 105^^^ clusters of barcodes were identified, suggesting that the barcode library has a diversity that is at least in the 105^^^ range.

#### Mice and *in utero* surgeries

All mouse colonies were maintained in accordance with protocols approved by the Bavarian government at the Max Planck Institute for Biological Intelligence or the Helmholtz-Zentrum München. C57BL/6 *wt* females were crossed to C57BL/6 *wt* or to CAS9-EGFP (B6.Gt(ROSA)26Sortm1.1(CAG-cas9*,-EGFP)Fezh/J, Jax 026179) males (Platt et al., 2014). Embryos were staged in days post-coitus, with E0.5 defined as 12:00 of a day that a vaginal plug was detected after overnight mating. Timed pregnant mice were anesthetized with isoflurane (5% induction, 2.5% during the surgery) and treated with the analgesic Metamizol (WDT). A microsyringe pump (Nanoject III Programmable Nano-liter Injector, 100/240V, DRUM3-000-207) was used to inject approximately 700 nl of DNA plasmid solution made of 0.6 *μ*g/*μ*l pEF1a-pBase (piggyBac-transposase; a gift from R. Platt) and the sgRNA plasmid 0.7 *μ*g/*μ*l, diluted in endo-free TE buffer and 0.002% Fast Green FCF (Sigma, F7252), into the lateral ventricle. pCAG-Cas9-EGFP (a gift from Randy Platt) plasmid was added when *wt* males were used for plugs. For TrackerSeq experiments, additionally barcode library (final concentarion 0.4 *μ*g/*μ*l) was added to DNA plasmid solution. Embryos were then electroporated by holding each head between large platinum-plated tweezer electrodes (5 mm in diameter, BTX, 45-0489) across the uterine wall, while 5 electric pulses (35 V, 50 ms at 1 Hz) were delivered with a square-wave electroporator (BTX, ECM830) (Saito, 2006). We used large electrodes in anticipation of targeting all areas of the GE (MGE, CGE, LGE) (Borrell et al., 2005). Pregnant dams were kept in single cages and pups were kept with their mothers. To assess cells distribution after *in utero* electroporation, embryos were collected after electroporation at E16.5 and E18.5. Dissected brains were fixed overnight in 4 % paraformaldehyde and after were washed with PBS. 50 *μ*m tissue sections were prepared on Leica VT1200S Vibratome and were mounted on slides with ProLong Glass Antifade Mountant (ThermoFisher). All images were acquired using STELLARIS 5 confocal microscope system (Leica). C57BL/6 wild type brains were prepared from E13.5 embryos, post-fixed in 4% PFA solution for 2.5 h and subsequently washed with PBS.

Before preparing brain tissue for scRNA-seq, each brain was examined under a stereomicroscope and only brains that met the following criteria were processed for scRNA-seq:

1. Dispersed tdTomato positive neurons throughout the neocortex. This indicates that we targeted MGE/CGE-derived INs that migrate tangentially from the ventral progenitors, where they are known to disperse to different cortical brain regions.
2. Dense tdTomato positive neurons throughout the striatum. MSNs are known to originate from the LGE and account for ca. 90% of the neurons in the striatum.
3. TdTomato positive neurons in the OB. GABAergic precursors are known to migrate from the LGE to the OB.

We performed immunohistochemical labelling to validate that after in IUE, individual brains express sgRNA in cortical IN derived from the MGE (anti-SST antibodies) and CGE (anti-Prox1) and in striatal MSNs derived from the LGE (anti-Ctip).

### Immunostainings

Paraformaldehyde-fixed brains at E13.5 and E18.5 were incubated in 10%/20%/30% sucrose for 24 h each, embedded in Neg-50™ Frozen Section Medium (Epredia) and subsequently snap-frozen in isobutane at -70°C. 16 *μ*m tissue sections were prepared on a Thermo Scientific CryoStar NX70 Cryostat and transferred to glass slides. Sections were incubated overnight with primary antibodies anti-MEIS2 (SCBT, sc-515470-AF594, 1:250), anti-LHX6 (SCBT, sc-271433-AF488, 1:50), anti-PROX1 (R&D Systems, AF2727, 1:250), anti-CTIP2 (Abcam, ab18465, 1:500). Sections were then incubated with secondary antibodies at room temperature for 2 h at 1:500 dilution: anti-rabbit AF594 (Invitrogen, A21207); anti-rat AF488 (Invitrogen, A21208); anti-goat AF488 (Invitrogen, A11055). Nuclei were counter-stained with DAPI and slides mounted with Aqua-Poly/Mount (Polysciences). Fluorescence imaging was conducted on a LSM880 confocal microscope (Zeiss Microscopy) using Plan-Apochromat 20/0.8 M27 or C-Aprochromat 63x/1.2 W Korr M27 objectives.

### Sample collection

We collected electroporated brains from mouse embryos at E16.5 in ice-cold Leibovitz’s L-15 Medium (ThermoFisher, 11415064) with 5% FBS or at P7-8 in ice-cold Hibernate-A Medium (ThermoFisher, A1247501) with 10% FBS and B-27 Supplement (ThermoFisher, 17504044). The same media were used during flow cytometry sorting. Only forebrains were collected, thus excluding the thalamus, hypothalamus, brainstem and cerebellum. Papain dissociation system (Wortington, LK003150) was carried out according to the protocol described in Jin et al., 2020 (Jin et al., 2020) on the gentleMACS™ Octo Dissociator (Miltenyi Biotec). To isolate positive cells, flow cytometry was done using a BD FACSAria III Cell Sorter (BD FACSDiva Software, version 8.0.2) with a 100-µm nozzle. EGFP and TdTomato-positive cells were collected in bulk for for testing sgRNA Meis2 knockout efficiency following *in vitro* protocol (above), or downstream processing on the 10x Genomics Chromium platform. After sorting 5,000–16,000 individual cells per sample, in PBS (Lonza) with 0.02% BSA (NEB), were loaded onto a 10X Genomics Chromium platform for Gel Beads-in-emulsion (GEM) and cDNA generation carrying cell- and transcript-specific barcode using the Chromium Single Cell 3’ Reagent Kit v3.1 with Feature Barcoding technology (PN-1000121) following manufacture protocol (document number CG000205, 10X Genomics).

### Logistic regression model to predict IN and PN genes

We used a recently published scRNA-seq data from (Bandler et al., 2022) to explore genes that are predictive for interneuron or projection neuron fate. Raw Counts for samples from GE-specific micro-dissections collected from WT mice at e13.5 and e15.5 were processed using Seurat (Hao et al., 2021). After integrating across batches, counts were normalized and scaled. Cluster annotations from (Bandler et al., 2022) were summarized into 4 broad cell classes: mitotic, trunk, interneuron and projection neuron. For performing logistic regression we subsetted cells from interneuron and projection neuron cell classes. Logistic regression was performed using 3000 most variable genes. To account for balanced design, cells were sub-sampled to have equal number of cells in both classes. A logistic regression model was trained on the scaled expression matrix of the corresponding cells and genes, where 2/3s of cells were used for training and the other third for validation. This was implemented using *cv*.*glmnet* (.., *family* = ”*binomial*”)-function from the R-package glmnet (Friedman et al., 2010). The model achieved 99.15% accuracy on the held-out validation set. For each gene, the model predicts a coefficient that reflects whether high expression of the gene is predictive of a cell being an interneuron (coefficient ∈ [0, 0.5]) or a projection neuron (coefficient ∈ [0.5, 1]).

### Preparation of tCROP-seq libraries

Uniquely barcoded RNA transcripts (poly(A)-mRNA and sgRNA) were reverse transcribed. 3’ Gene Expression library and CRISPR Screening library were generated according to manufacturer’s user guide (Document number CG000205) with use of Chromium Library v3.1 kit (PN-1000121), Feature Barcode Library Kit (PN-1000079) and Single Index Kit (PN-1000213) (10X Genomics). Libraries were quantified with Agilent BioAnalyzer.

### Preparation of TrackerSeq NGS libraries

The TrackerSeq lineage libraries retrieved from cDNA were amplified with Q5 polymerase (NEB, M094S) in a 50-µl reaction, using 10 µl of cDNA as template (Bandler et al., 2022). Specifically, each PCR contained: 25 µl Q5 High-fidelity 2X Master Mix, 2.5 µl 10 µM P7-indexed reverse primer, 2.5 µl 10 µM i5-indexed forward primer, 10 µl molecular grade H_2_O, 10 µl cDNA (for primer sequences and indices, see Table S1). Libraries were purified with a dual-sided selection using SPRIselect (Beckman Coulter, B23318), and quantified with an Agilent BioAnalyzer.

### Sequencing and read mapping

Transcriptome and CRISPR barcode libraries were sequenced either on an Illumina NextSeq 500 at the Next Generation Sequencing Facility of the Max Planck Institute of Biochemistry or on a NovaSeq at the Genomics Core Facility at the Helmholtz Center in Munich. For a detailed report on each dataset, see Table S9. Sequencing reads in FASTQ files were aligned to a reference transcriptome (mm10-2.1.0) and collapsed into UMI counts using the 10x Genomics Cell Ranger software (version 3.0.2 or 5.0.1).

### tCROP-seq pre-processing

UMI count data was loaded into R and processed using the Seurat v4 package (Hao et al., 2021). CRISPR gRNAs were recovered using CellRanger (Zheng et al., 2017), which produces an output CSV file containing the cell barcodes and the sgRNA detected in that cell.

#### Processing embryonic tCROP-seq datasets

Electroporation of ventral progenitors using the 5 mm electrode targets some additional progenitors located adjacent to the ganglionic eminence. These include progenitors of excitatory neurons located at the border between the pallium and the subpallium. Thus, our data set consisted of: Inhibitory: 16098 neurons; Excitatory: 10010 neurons; Glial: 5915 cells; Pericytes: 1008 cells; Fibroblasts: 537 cells; Macrophages: 523 cells; Blood: 390 cells; We focused only on cells from inhibitory neurons and excluded the others. We integrated inhibitory neurons with scRNA-seq datasets from wild-type mice (Bandler et al., 2022) to get a higher resolution of inhibitory cell states (Figure1b) using the integration tool from Seurat (Hao et al., 2021). We obtained cluster-specific marker genes by performing differential expression analysis (see below). Clusters were assigned to cell types based on the expression of known marker genes, primarily using http://mousebrain.org/development/ (La Manno et al., 2021) and https://DropViz.org (Saunders et al., 2018).

#### Processing postnatal tCROP-seq datasets

To process the P7 datasets, we integrated Harmony (v1.0, (Korsunsky et al., 2019)) into our Seurat (Hao et al., 2021) workflow for batch correction, using default settings (theta = 2, lambda = 1, sigma = 0.1). We used the first 30 Harmony embeddings for UMAP (https://github.com/lmcinnes/umap) visualizations and clustering analysis. To group cells into clusters, we first constructed a shared-nearest neighbour graph from Harmony embeddings using the FindNeighbors() algorithm, then input the graph into the FindClusters() function in Seurat (dimensions = 30, res = 0.8). To test whether our postnatal dataset was subject to non-specific background expression, we applied DecontX (Yang et al., 2020) using the default parameters. We retrieved the count matrix from our Seurat object, created an SCE object, ran DecontX and then added the corrected count matrix back to the Seurat object. The differences before and after correction are relatively small. Therefore, we decided to use the uncorrected counts for the subsequent analysis.

### Comparing cell type composition between perturbations

We compared the perturbation effect on cell type composition using the method described by Jin et al. (Jin et al., 2020). A detailed script of the analysis is deposited on a https://github.com/mayer-lab/Dvoretskova-et-al. Compositional change was investigated using the “CellComp_Poisson” R function from Jin et al., 2020 (Jin et al., 2020). It performs Poisson regression analysis to identify genes that are differentially expressed across different cell types, perturbations and batches. The function first performs data cleaning by creating a metadata data frame and filtering out cells with low counts. It then fits a Poisson regression model for each combination of cell type and perturbation and extracts the coefficients for the perturbation variable. These coefficients are then used to calculate p-values and adjusted p-values for each gene.

### Differential gene expression analysis

We used the Libra package to perform differential gene expression analysis (Squair et al., 2021). We ran the run_DE function on Seurat objects using the following parameters: de_family = pseudobulk, de_family = pseudobulk, de_method = edgeR, de_type = LRT. We obtained DEGs of PNs or INs by using run_DE function on cells grouped into classes (mitotic, projection neurons, and interneurons). We filtered for statistically significant genes (FDR-adjusted p-value threshold = 0.05). Genes were considered differentially expressed if avg_logFC < -1.0 or avg_logFC > 1.0.

We also utilized the R packagage “Libra” to calculate the differentially expressed (DE) genes for each cluster (i_Calb2/Nxph1, i_Cck/Reln, i_Ebf1/Zfp503, i_Foxp1/Isl1, i_Foxp1/Six3, i_Isl1/Bcl11b, i_Lhx6/Npy, i_Meis2/Bcl11b, i_Nfib/Tcf4, i_Nr2f2/Nnat, i_Tiam2/Zfp704, i_Tshz1/Pbx1). The result of the DE analysis is in Table S4 (See attached table taken from our submission). We applied the threshold *p*_*val*_*adj*_ < 0.05&(*avg*_*logFC* < −1.0|*avg*_*logFC* > 1.0) to select the genes for intersection with the Chip-seq data. We combined DE genes from all subtypes, and in up or down regulated genes we took unique gene symbols for the Venn diagram.

### TrackerSeq (lineage tracing) barcode processing and analysis

For a subset of datasets (ED210204, ED210215, ED211111, ED211124), we included TrackerSeq lineage barcodes to perform a clonal analysis. We followed the protocol outlined in (Bandler et al., 2022) to process the TrackerSeq barcodes and obtain cloneIDs for each corresponding cell barcode. The resulting cloneIDs were added to the Seurat object metadata. To quantify clonal relationship between cell classes, the inhibitory clusters were first merged into cell classes (Figure 2c) based on whether they were annotated as mitotic (*Ube2c* and *Top2a*), or as INs and PNs (*Gad2*). The UpsetR library was used to count the number of clones shared between the neuronal classes, as well as the proportion of clonal relationships in gMeis2 and gLacZ datasets. The set size is the number of cells in the class. The UpSet bar plot shows the calculated proportion of each type of clonal distribution category within the perturbation. Each percentage was obtained by dividing the clones belonging to that category (e.g. clones containing only mitotic and INs) by the number of clones belonging to all other categories of clonal distribution.

To quantify lineage coupling, we used a method from Weinreb et al. 2020 (Weinreb et al., 2020). The method computes an observed/expected ratio of shared barcodes for each pair of cell-states. A barcode is considered shared if it appears in at least one cell from both states. From the observed shared barcode matrix *O*_*i* *j*_, it derives an expected shared barcode matrix *E*_*i* *j*_ under the assumption of no lineage couplings, as follows:

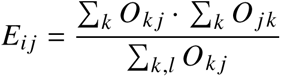

To avoid artifacts from particularly large or atypical clones, it re-computed these matrices 1000 times, each time using a random 25% sample of clones. The lineage coupling scores shown in Figure S2g represent the median *O*_*i* *j*_ /*E*_*i* *j*_ from these 1000 randomized trials.

### Hotspot analysis of gene coexpression

Hotspot(v0.91) is a tool for identifying co-expressing gene modules in a single-cell dataset (DeTomaso and Yosef, 2021). It computes gene modules by evaluating the pairwise correlation of genes with high local autocorrelation, then clusters the results into a gene-gene affinity matrix. The *Gad2*-expressing inhibitory neuron population in the P7 dataset was first subset from the remainder of the dataset to identify inhibitory specific modules in the embryonic dataset. We ran the depth-adjusted negative binomial model on the entire count matrix and Harmony (v1.0) corrected principal components. We computed a k-nearest-neighbors (KNN) graph with 30 neighbors, 9154 non-varying genes were subsequently detected and removed. Autocorrelations between each gene were calculated, and the top 500 significant (FDR < 0.05) genes were used to evaluate pairwise gene associations (local correlations). After pairwise local correlations were calculated, we grouped genes into modules. Modules were created through agglomerative clustering, where the minimum number of genes per module was set to 30. 8 modules were identified, and 103 genes were not assigned to a module. Summary per-cell module scores are calculated using the calculate_module_scores() function as described by DeTomaso et al. (DeTomaso and Yosef, 2021). As described by Jin et al. 2020, linear regression was used to test the relationship between perturbation and Hotspot module gene scores (Jin et al., 2020).

### GO Term analysis of differentially expressed genes and module genes

GO Term analysis was done using the package enrichR (Kuleshov et al., 2016). The DEGs and module genes of each module were queried against the following databases: GO_Molecular_Functio_2018, GO_Cellular_Component_2018, and GO_Biological_Process_2018. Only GO Terms that were significant (p-value adjusted < 0.05) were kept.

### Luciferase assay

Regulatory elements were amplified from mouse genomic DNA with Q5 polymerase (NEB, M0491) using primers listed in Table S8 and cloned into pGL4.24[luc2P/minP] (Promega, E8421) with NEBuilder HiFi DNA Assembly kit (NEB, E2621). Enhancer hs1080 had to be cloned in reverse-complement. Mouse *Meis2* isoform D (4) (the tag was removed) and *Lhx6* variant 1 (C-DYK) expressing vectors were purchased from Genscript, *Dlx5* and *Pbx1* coding sequences were amplified from mouse cDNA and cloned into pcDNA3.1 (Genscript). Meis2 vector was mutated with NEBuilder HiFi DNA Assembly kit (NEB, E2621) to harbor the human mutation p.(Arg333Lys), c.998G>A (Meis2*333) (Verheije et al., 2019). A short version of enhD1 luciferase vector was mutated with use of a gBlock (IDT) and NEBuilder HiFi DNA Assembly kit (NEB, E2621). Luciferase reporter vectors were co-transfected with pNL1.1.PGK[Nluc/PGK] (N1441, Promega), pcDNA3 or pcDNA3-Dlx5, pcDNA3-PBX1, pcDNA3-Meis2, pcDNA3-Lhx6. Neuro2a cells were seeded in 24-well plates at 80,000 cells per well and on the next day were transfected with TransIT-LT1 Transfection Reagent (Mirus, MIR 2300), using 150 ng luciferase reporter, 10 ng Nluc/PGK and 350 ng total of pcDNA3.1 plasmids per well (150 ng per TFs vector). pcDNA stands for a control plasmid pcDNA3.1 which does not contain a protein coding sequence. The pcDNA was used to balance DNA load during transfections. Cells were harvested 24 hours after transfection and luciferases activity was measured using Nano-Glo® Dual-Luciferase® Reporter Assay System (Promega) on Berthold Multimode reader Tristar2S. A Nanoluc reporter was used for normalization. Statistical tests were performed using the GraphPad Prism software (Version 10.0.2). Two-way analysis of variance (ANOVA) followed by Tukey’s honestly significant difference (HSD) test were used to determine the statistical significance between various conditions. All results for statistical analysis are listed in Table S8.

### Chromatin-Immunoprecipitation (ChIP)

Mice were handled in accordance with the CNIC Ethics Committee, Spanish laws, and the EU Directive 2010/63/EU for the use of animals in research. GEs and part of the underlying striatum of 70 *wt* C57BL/6 embryos at E14.5 were microdissected and immediately fixed in 1% formaldehyde for 5 min. Tissue preparation, immunoprecipitation and sequencing on an Illumina HiSeq2500 were performed as previously described (Delgado et al., 2021). Immunoprecipitation was carried out using a combination of two anti-MEIS antibodies, one recognizing MEIS1A and MEIS2A, the other recognizing all MEIS2 isoforms (Mercader et al., 2005).

### ChIP-seq Data Analysis

61 bp single-end reads were trimmed using Cutadapt (v1.16) and mapped to GRCm38 using Bowtie2 (v2.3.0) (Langmead and Salzberg, 2012) followed by peak calling with MACS2 (v2.1.2) (Feng et al., 2012) using a cutoff of q=0.01. TSS definitions were adapted from the eukaryotic promoter database (mmEPDnew version 003) (Meylan et al., 2020). We determined the distance of each peak to the nearest TSS using the R package plyranges (v1.180). Using custom R scripts, peaks were assigned to the TSS of a gene when overlapping a ca. 5 kb region around a TSS, defined as promoter region. Overlap with developmental enhancers (Gorkin et al., 2020) was determined in the same way. Similarly, we determined overlap of MEIS2 binding sites with DLX5 binding sites at E13.5 from Lindtner et al. (Lindtner et al., 2019) and LHX6 binding sites at E13.5 from Sandberg et al. (Sandberg et al., 2016). Enrichment of enhancer-overlapping peaks among shared MEIS2/DLX5 peaks, compared to MEIS2- and DLX5-exclusive peaks, was determined using Peason’s Chi-squared test of the R ‘stats’ package(v4.0.2). Genomic tracks and Vista enhancers (Visel et al., 2007) were visualized using the Integrated Genomics Viewer (v2.4.1) (Robinson et al., 2011).

Motif identification and enrichment of known motifs were carried out by HOMER (v4.10.4) (Heinz et al., 2010) using default settings. Motif enrichment within enhancer- and promoter-overlapping peaks was likewise performed with HOMER. We used SpaMo (v5.4.1) (Whitington et al., 2011)) to determine motif spacing between MEIS2 and DLX5 binding motifs in common MEIS2/DLX5 binding sites, within 100 bp up- and down-stream of MEIS2 peak summits.

### Data used in this study

GSE167047 (snATAC-seq of E12.5 MGE and LGE (Rhodes et al., 2022)), GSE85705 (LHX6-ChIP-seq GE E13.5 (Sandberg et al., 2016)), GSE124936 (DLX1, DLX2 & DLX5-ChIP-seq GE E13.5 (Lindtner et al., 2019)) and GSE188528 (scRNA-seq of LGE, MGE, CGE E13.5 (Bandler et al., 2022)) were downloaded from GEO. Developmental enhancers and interacting genes (Gorkin et al., 2020). Vista enhancer images were downloaded from the Vista Enhancer Browser (Visel et al., 2007). ChIP-seq and ATAC-seq tracks were presented using the IGV software (Thorvaldsdóttir et al., 2013)

## Code availability

The analyses described here are available on github.com/mayer-lab

## References

Agoston, Z., Heine, P., Brill, M. S., Grebbin, B. M., Hau, A.-C., Kallenborn-Gerhardt, W., Schramm, J., Götz, M., and Schulte, D. (2014). Meis2 is a pax6 co-factor in neurogenesis and dopaminergic periglomerular fate specification in the adult olfactory bulb. Development, 141(1):28–38.

Allaway, K. C. and Machold, R. (2017). Developmental specification of forebrain cholinergic neurons. Dev Biol, 421(1):1–7.

Anderson, S. A., Eisenstat, D. D., Shi, L., and Rubenstein, J. L. (1997). Interneuron migration from basal forebrain to neocortex: dependence on dlx genes. Science, 278(5337):474–6.

Anderson, S. A., Marín, O., Horn, C., Jennings, K., and Rubenstein, J. L. (2001). Distinct cortical migrations from the medial and lateral ganglionic eminences. Development, 128(3):353–63.

Asgarian, Z., Oliveira, M. G., Stryjewska, A., Maragkos, I., Rubin, A. N., Magno, L., Pachnis, V., Ghorbani, M., Hiebert, S. W., Denaxa, M., and Kessaris, N. (2022). Mtg8 interacts with lhx6 to specify cortical interneuron subtype identity. Nat Commun, 13(1):5217.

Bandler, R. C., Mayer, C., and Fishell, G. (2017). Cortical interneuron specification: the juncture of genes, time and geometry. Curr Opin Neurobiol, 42:17–24.

Bandler, R. C., Vitali, I., Delgado, R. N., Ho, M. C., Dvoretskova, E., Ibarra Molinas, J. S., Frazel, P. W., Mohammadkhani, M., Machold, R., Maedler, S., Liddelow, S. A., Nowakowski, T. J., Fishell, G., and Mayer, C. (2022). Single-cell delineation of lineage and genetic identity in the mouse brain. Nature, 601(7893):404–409.

Batista-Brito, R., Machold, R., Klein, C., and Fishell, G. (2008). Gene expression in cortical interneuron precursors is prescient of their mature function. Cereb Cortex, 18(10):2306–17.

Berenguer, M. and Duester, G. (2022). Retinoic acid, rars and early development. J Mol Endocrinol, 69(4):T59–T67.

Bonev, B., Mendelson Cohen, N., Szabo, Q., Fritsch, L., Papadopoulos, G. L., Lubling, Y., Xu, X., Lv, X., Hugnot, J.-P., Tanay, A., and Cavalli, G. (2017). Multiscale 3d genome rewiring during mouse neural development. Cell, 171(3):557–572.e24.

Borello, U., Cobos, I., Long, J. E., McWhirter, J. R., Murre, C., and Rubenstein, J. L. R. (2008). Fgf15 promotes neurogenesis and opposes fgf8 function during neocortical development. Neural Dev, 3:17.

Borrell, V., Yoshimura, Y., and Callaway, E. M. (2005). Targeted gene delivery to telencephalic inhibitory neurons by directional in utero electroporation. J Neurosci Methods, 143(2):151–8.

Bridoux, L., Zarrineh, P., Mallen, J., Phuycharoen, M., Latorre, V., Ladam, F., Losa, M., Baker, S. M., Sagerstrom, C., Mace, K. A., Rattray, M., and Bobola, N. (2020). Hox paralogs selectively convert binding of ubiquitous transcription factors into tissue-specific patterns of enhancer activation. PLoS Genet, 16(12):e1009162.

Butt, S. J. B., Fuccillo, M., Nery, S., Noctor, S., Kriegstein, A., Corbin, J. G., and Fishell, G. (2005). The temporal and spatial origins of cortical interneurons predict their physiological subtype. Neuron, 48(4):591–604.

Cesario, J. M., Landin Malt, A., Deacon, L. J., Sandberg, M., Vogt, D., Tang, Z., Zhao, Y., Brown, S., Rubenstein, J. L., and Jeong, J. (2015). Lhx6 and lhx8 promote palate development through negative regulation of a cell cycle inhibitor gene, p57kip2. Hum Mol Genet, 24(17):5024–39.

Chang, C. P., Jacobs, Y., Nakamura, T., Jenkins, N. A., Copeland, N. G., and Cleary, M. L. (1997). Meis proteins are major in vivo dna binding partners for wild-type but not chimeric pbx proteins. Mol Cell Biol, 17(10):5679–87.

Chapman, H., Riesenberg, A., Ehrman, L. A., Kohli, V., Nardini, D., Nakafuku, M., Campbell, K., and Waclaw, R. R. (2018). Gsx transcription factors control neuronal versus glial specification in ventricular zone progenitors of the mouse lateral ganglionic eminence. Dev Biol, 442(1):115–126.

Chatzi, C., Brade, T., and Duester, G. (2011). Retinoic acid functions as a key gabaergic differentiation signal in the basal ganglia. PLoS Biol, 9(4):e1000609.

Chenn, A. and Walsh, C. A. (2002). Regulation of cerebral cortical size by control of cell cycle exit in neural precursors. Science, 297(5580):365–9.

Datlinger, P., Rendeiro, A. F., Schmidl, C., Krausgruber, T., Traxler, P., Klughammer, J., Schuster, L. C., Kuchler, A., Alpar, D., and Bock, C. (2017). Pooled crispr screening with single-cell transcriptome readout. Nat Methods, 14(3):297–301.

Delás, M. J., Kalaitzis, C. M., Fawzi, T., Demuth, M., Zhang, I., Stuart, H. T., Costantini, E., Ivanovitch, K., Tanaka, E. M., and Briscoe, J. (2023). Developmental cell fate choice in neural tube progenitors employs two distinct cis-regulatory strategies. Dev Cell, 58(1):3–17.e8.

Delgado, I., Giovinazzo, G., Temiño, S., Gauthier, Y., Balsalobre, A., Drouin, J., and Torres, M. (2021). Control of mouse limb initiation and antero-posterior patterning by meis transcription factors. Nat Commun, 12(1):3086.

den Hoed, J., Devaraju, >K., and Fisher, S. E. (2021). Molecular networks of the foxp2 transcription factor in the brain. EMBO Rep, 22(8):e52803.

DeTomaso, D. and Yosef, N. (2021). Hotspot identifies informative gene modules across modalities of single-cell genomics. Cell Syst, 12(5):446–456.e9.

Ding, S., Wu, X., Li, G., Han, M., Zhuang, Y., and Xu, T. (2005). Efficient transposition of the piggybac (pb) transposon in mammalian cells and mice. Cell, 122(3):473–83.

Dodson, P. D., Larvin, J. T., Duffell, J. M., Garas, F. N., Doig, N. M., Kessaris, N., Duguid, I. C., Bogacz, R., Butt, S. J. B., and Magill, P. J. (2015). Distinct developmental origins manifest in the specialized encoding of movement by adult neurons of the external globus pallidus. Neuron, 86(2):501–13.

Doench, J. G., Fusi, N., Sullender, M., Hegde, M., Vaimberg, E. W., Donovan, K. F., Smith, I., Tothova, Z., Wilen, C., Orchard, R., Virgin, H. W., Listgarten, J., and Root, D. E. (2016). Optimized sgrna design to maximize activity and minimize off-target effects of crispr-cas9. Nat Biotechnol, 34(2):184–191.

Douglas, G., Cho, M. T., Telegrafi, A., Winter, S., Carmichael, J., Zackai, E. H., Deardorff, M. A., Harr, M., Williams, L., Psychogios, A., Erwin, A. L., Grebe, T., Retterer, K., and Juusola, J. (2018). De novo missense variants in meis2 recapitulate the microdeletion phenotype of cardiac and palate abnormalities, developmental delay, intellectual disability and dysmorphic features. Am J Med Genet A, 176(9):1845–1851.

Dupacova, N., Antosova, B., Paces, J., and Kozmik, Z. (2021). Meis homeobox genes control progenitor competence in the retina. Proc Natl Acad Sci U S A, 118(12).

Feng, J., Liu, T., Qin, B., Zhang, Y., and Liu, X. S. (2012). Identifying chip-seq enrichment using macs. Nat Protoc, 7(9):1728–40.

Fischer, E. S., Böhm, K., Lydeard, J. R., Yang, H., Stadler, M. B., Cavadini, S., Nagel, J., Serluca, F., Acker, V., Lingaraju, G. M., Tichkule, R. B., Schebesta, M., Forrester, W. C., Schirle, M., Hassiepen, U., Ottl, J., Hild, M., Beckwith, R. E. J., Harper, J. W., Jenkins, J. L., and Thomä, N. H. (2014). Structure of the ddb1-crbn e3 ubiquitin ligase in complex with thalidomide. Nature, 512(7512):49–53.

Flames, N., Pla, R., Gelman, D., Rubenstein, J., Puelles, L., and Marín, O. (2007). Delineation of multiple subpallial progenitor domains by the combinatorial expression of transcriptional codes. J. Neurosci.; Journal of Neuroscience, 27(36):9682–9695.

French, C. A. and Fisher, S. E. (2014). What can mice tell us about foxp2 function? Curr Opin Neurobiol, 28:72–9.

Friedman, J., Hastie, T., and Tibshirani, R. (2010). Regularization paths for generalized linear models via coordinate descent. J Stat Softw, 33(1):1–22.

Gangfuß, A., Yigit, G., Altmüller, J., Nürnberg, P., Czeschik, J. C., Wollnik, B., Bögershausen, N., Burfeind, P., Wieczorek, D., Kaiser, F., Roos, A., Kölbel, H., Schara-Schmidt, U., and Kuechler, A. (2021). Intellectual disability associated with craniofacial dysmorphism, cleft palate, and congenital heart defect due to a de novo meis2 mutation: A clinical longitudinal study. Am J Med Genet A, 185(4):1216–1221.

Gelman, D., Griveau, A., Dehorter, N., Teissier, A., Varela, C., Pla, R., Pierani, A., and Marín, O. (2011). A wide diversity of cortical gabaergic interneurons derives from the embryonic preoptic area. J Neurosci, 31(46):16570–80.

Gerfen, C. R. and Surmeier, D. J. (2011). Modulation of striatal projection systems by dopamine. Annu Rev Neurosci, 34:441–66.

Giliberti, A., Currò, A., Papa, F. T., Frullanti, E., Ariani, F., Coriolani, G., Grosso, S., Renieri, A., and Mari, F. (2020). Meis2 gene is responsible for intellectual disability, cardiac defects and a distinct facial phenotype. Eur J Med Genet, 63(1):103627.

Gorkin, D. U., Barozzi, I., Zhao, Y., Zhang, Y., Huang, H., Lee, A. Y., Li, B., Chiou, J., Wildberg, A., Ding, B., Zhang, B., Wang, M., Strattan, J. S., Davidson, J. M., Qiu, Y., Afzal, V., Akiyama, J. A., Plajzer-Frick, I., Novak, C. S., Kato, M., Garvin, T. H., Pham, Q. T., Harrington, A. N., Mannion, B. J., Lee, E. A., Fukuda-Yuzawa, Y., He, Y., Preissl, S., Chee, S., Han, J. Y., Williams, B. A., Trout, D., Amrhein, H., Yang, H., Cherry, J. M., Wang, W., Gaulton, K., Ecker, J. R., Shen, Y., Dickel, D. E., Visel, A., Pennacchio, L. A., and Ren, B. (2020). An atlas of dynamic chromatin landscapes in mouse fetal development. Nature, 583(7818):744–751.

Hao, Y., Hao, S., Andersen-Nissen, E., Mauck, 3rd, W. M., Zheng, S., Butler, A., Lee, M. J., Wilk, A. J., Darby, C., Zager, M., Hoffman, P., Stoeckius, M., Papalexi, E., Mimitou, E. P., Jain, J., Srivastava, A., Stuart, T., Fleming, L. M., Yeung, B., Rogers, A. J., McElrath, J. M., Blish, C. A., Gottardo, R., Smibert, P., and Satija, R. (2021). Integrated analysis of multimodal single-cell data. Cell, 184(13):3573–3587.e29.

Heinz, S., Benner, C., Spann, N., Bertolino, E., Lin, Y. C., Laslo, P., Cheng, J. X., Murre, C., Singh, H., and Glass, C. K. (2010). Simple combinations of lineage-determining transcription factors prime cis-regulatory elements required for macrophage and b cell identities. Mol Cell, 38(4):576–89.

Hobert, O. and Westphal, H. (2000). Functions of lim-homeobox genes. Trends Genet, 16(2):75– 83.

Hsiau, T., Conant, D., Rossi, N., Maures, T., Waite, K., Yang, J., Joshi, S., Kelso, R., Holden, K., Enzmann, B., and Stoner, R. (2019). Inference of crispr edits from sanger trace data. bioRxiv.

Hyman-Walsh, C., Bjerke, G. A., and Wotton, D. (2010). An autoinhibitory effect of the homothorax domain of meis2. FEBS J, 277(12):2584–97.

Jin, X., Simmons, S. K., Guo, A., Shetty, A. S., Ko, M., Nguyen, L., Jokhi, V., Robinson, E., Oyler, P., Curry, N., Deangeli, G., Lodato, S., Levin, J. Z., Regev, A., Zhang, F., and Arlotta, P. (2020). In vivo perturb-seq reveals neuronal and glial abnormalities associated with autism risk genes. Science, 370(6520).

Jindal, G. A. and Farley, E. K. (2021). Enhancer grammar in development, evolution, and disease: dependencies and interplay. Dev Cell, 56(5):575–587.

Jolma, A., Yin, Y., Nitta, K. R., Dave, K., Popov, A., Taipale, M., Enge, M., Kivioja, T., Morgunova, E., and Taipale, J. (2015). Dna-dependent formation of transcription factor pairs alters their binding specificity. Nature, 527(7578):384–8.

Kim, H., Berens, N. C., Ochandarena, N. E., and Philpot, B. D. (2020). Region and cell type distribution of tcf4 in the postnatal mouse brain. Front Neuroanat, 14:42.

Kim, S., Morgunova, E., Naqvi, S., Bader, M., Koska, M., Popov, A., Luong, C., Pogson, A., Claes, P., Taipale, J., and Wysocka, J. (2023). Dna-guided transcription factor cooperativity shapes face and limb mesenchyme. bioRxiv.

Knowles, R., Dehorter, N., and Ellender, T. (2021). From progenitors to progeny: Shaping striatal circuit development and function. J Neurosci, 41(46):9483–9502.

Ko, H.-A., Chen, S.-Y., Chen, H.-Y., Hao, H.-J., and Liu, F.-C. (2013). Cell type-selective expression of the zinc finger-containing gene nolz-1/zfp503 in the developing mouse striatum. Neurosci Lett, 548:44–9.

Korsunsky, I., Millard, N., Fan, J., Slowikowski, K., Zhang, F., Wei, K., Baglaenko, Y., Brenner, M., Loh, P.-R., and Raychaudhuri, S. (2019). Fast, sensitive and accurate integration of single-cell data with harmony. Nat Methods, 16(12):1289–1296.

Kreitzer, A. C. and Malenka, R. C. (2008). Striatal plasticity and basal ganglia circuit function. Neuron, 60(4):543–54.

Kuleshov, M. V., Jones, M. R., Rouillard, A. D., Fernandez, N. F., Duan, Q., Wang, Z., Koplev, S., Jenkins, S. L., Jagodnik, K. M., Lachmann, A., McDermott, M. G., Monteiro, C. D., Gundersen, G. W., and Ma’ayan, A. (2016). Enrichr: a comprehensive gene set enrichment analysis web server 2016 update. Nucleic Acids Res, 44(W1):W90–7.

La Manno, G., Siletti, K., Furlan, A., Gyllborg, D., Vinsland, E., Mossi Albiach, A., Mattsson Langseth, C., Khven, I., Lederer, A. R., Dratva, L. M., Johnsson, A., Nilsson, M., Lönnerberg, P., and Linnarsson, S. (2021). Molecular architecture of the developing mouse brain. Nature, 596(7870):92–96.

Langmead, B. and Salzberg, S. L. (2012). Fast gapped-read alignment with bowtie 2. Nat Methods, 9(4):357–9.

Lee, D. R., Rhodes, C., Mitra, A., Zhang, Y., Maric, D., Dale, R. K., and Petros, T. J. (2022). Transcriptional heterogeneity of ventricular zone cells in the ganglionic eminences of the mouse forebrain. Elife, 11.

Leung, R. F., George, A. M., Roussel, E. M., Faux, M. C., Wigle, J. T., and Eisenstat, D. D. (2022). Genetic regulation of vertebrate forebrain development by homeobox genes. Front Neurosci, 16:843794.

Lim, L., Mi, D., Llorca, A., and Marín, O. (2018). Development and functional diversification of cortical interneurons. Neuron, 100(2):294–313.

Lindtner, S., Catta-Preta, R., Tian, H., Su-Feher, L., Price, J. D., Dickel, D. E., Greiner, V., Silberberg, S. N., McKinsey, G. L., McManus, M. T., Pennacchio, L. A., Visel, A., Nord, A. S., and Rubenstein, J. L. R. (2019). Genomic resolution of dlx-orchestrated transcriptional circuits driving development of forebrain gabaergic neurons. Cell Rep, 28(8):2048–2063.e8.

Longobardi, E., Penkov, D., Mateos, D., De Florian, G., Torres, M., and Blasi, F. (2014). Biochemistry of the tale transcription factors prep, meis, and pbx in vertebrates. Dev Dyn, 243(1):59–75.

Louw, J. J., Corveleyn, A., Jia, Y., Hens, G., Gewillig, M., and Devriendt, K. (2015). Meis2 involvement in cardiac development, cleft palate, and intellectual disability. Am J Med Genet A, 167A(5):1142–6.

Mayer, C., Hafemeister, C., Bandler, R. C., Machold, R., Batista Brito, R., Jaglin, X., Allaway, K., Butler, A., Fishell, G., and Satija, R. (2018). Developmental diversification of cortical inhibitory interneurons. Nature, 555(7697):457–462.

Megason, S. G. and McMahon, A. P. (2002). A mitogen gradient of dorsal midline wnts organizes growth in the cns. Development, 129(9):2087–98.

Mercader, N., Tanaka, E. M., and Torres, M. (2005). Proximodistal identity during vertebrate limb regeneration is regulated by meis homeodomain proteins. Development, 132(18):4131–42.

Mesman, S., Bakker, R., and Smidt, M. P. (2020). Tcf4 is required for correct brain development during embryogenesis. Molecular and Cellular Neuroscience, 106:103502.

Meylan, P., Dreos, R., Ambrosini, G., Groux, R., and Bucher, P. (2020). Epd in 2020: enhanced data visualization and extension to ncrna promoters. Nucleic Acids Res, 48(D1):D65–D69.

Miyoshi, G., Hjerling-Leffler, J., Karayannis, T., Sousa, V. H., Butt, S. J. B., Battiste, J., Johnson, J. E., Machold, R. P., and Fishell, G. (2010). Genetic fate mapping reveals that the caudal ganglionic eminence produces a large and diverse population of superficial cortical interneurons. J Neurosci, 30(5):1582–94.

Miyoshi, G., Young, A., Petros, T., Karayannis, T., McKenzie Chang, M., Lavado, A., Iwano, T., Nakajima, M., Taniguchi, H., Huang, Z. J., Heintz, N., Oliver, G., Matsuzaki, F., Machold, R. P., and Fishell, G. (2015). Prox1 regulates the subtype-specific development of caudal ganglionic eminence-derived gabaergic cortical interneurons. J Neurosci, 35(37):12869–89.

Molotkova, N., Molotkov, A., and Duester, G. (2007). Role of retinoic acid during forebrain development begins late when raldh3 generates retinoic acid in the ventral subventricular zone. Dev Biol, 303(2):601–10.

Nery, S., Fishell, G., and Corbin, J. G. (2002). The caudal ganglionic eminence is a source of distinct cortical and subcortical cell populations. Nat Neurosci, 5(12):1279–87.

Ng, F. S. L., Schütte, J., Ruau, D., Diamanti, E., Hannah, R., Kinston, S. J., and Göttgens, B. (2014). Constrained transcription factor spacing is prevalent and important for transcriptional control of mouse blood cells. Nucleic Acids Res, 42(22):13513–24.

Panganiban, G. and Rubenstein, J. L. R. (2002). Developmental functions of the distal-less/dlx homeobox genes. Development, 129(19):4371–86.

Platt, R. J., Chen, S., Zhou, Y., Yim, M. J., Swiech, L., Kempton, H. R., Dahlman, J. E., Parnas, O., Eisenhaure, T. M., Jovanovic, M., Graham, D. B., Jhunjhunwala, S., Heidenreich, M., Xavier, R. J., Langer, R., Anderson, D. G., Hacohen, N., Regev, A., Feng, G., Sharp, P. A., and Zhang, F. (2014). Crispr-cas9 knockin mice for genome editing and cancer modeling. Cell, 159(2):440–55.

Replogle, J. M., Norman, T. M., Xu, A., Hussmann, J. A., Chen, J., Cogan, J. Z., Meer, E. J., Terry, J. M., Riordan, D. P., Srinivas, N., Fiddes, I. T., Arthur, J. G., Alvarado, L. J., Pfeiffer, K. A., Mikkelsen, T. S., Weissman, J. S., and Adamson, B. (2020). Combinatorial single-cell crispr screens by direct guide rna capture and targeted sequencing. Nat Biotechnol, 38(8):954–961.

Rhodes, C. T., Thompson, J. J., Mitra, A., Asokumar, D., Lee, D. R., Lee, D. J., Zhang, Y., Jason, E., Dale, R. K., Rocha, P. P., and Petros, T. J. (2022). An epigenome atlas of neural progenitors within the embryonic mouse forebrain. Nat Commun, 13(1):4196.

Robinson, J. T., Thorvaldsdóttir, H., Winckler, W., Guttman, M., Lander, E. S., Getz, G., and Mesirov, J. P. (2011). Integrative genomics viewer. Nat Biotechnol, 29(1):24–6.

Saito, T. (2006). In vivo electroporation in the embryonic mouse central nervous system. Nat Protoc, 1(3):1552–8.

Sandberg, M., Flandin, P., Silberberg, S., Su-Feher, L., Price, J. D., Hu, J. S., Kim, C., Visel, A., Nord, A. S., and Rubenstein, J. L. R. (2016). Transcriptional networks controlled by nkx2-1 in the development of forebrain gabaergic neurons. Neuron, 91(6):1260–1275.

Sandberg, M., Taher, L., Hu, J., Black, B. L., Nord, A. S., and Rubenstein, J. L. R. (2018). Genomic analysis of transcriptional networks directing progression of cell states during mge development. Neural Dev, 13(1):21.

Sanson, K. R., Hanna, R. E., Hegde, M., Donovan, K. F., Strand, C., Sullender, M. E., Vaimberg, E. W., Goodale, A., Root, D. E., Piccioni, F., and Doench, J. G. (2018). Optimized libraries for crispr-cas9 genetic screens with multiple modalities. Nat Commun, 9(1):5416.

Saunders, A., Macosko, E. Z., Wysoker, A., Goldman, M., Krienen, F. M., de Rivera, H., Bien, E., Baum, M., Bortolin, L., Wang, S., Goeva, A., Nemesh, J., Kamitaki, N., Brumbaugh, S., Kulp, D., and McCarroll, S. A. (2018). Molecular diversity and specializations among the cells of the adult mouse brain. Cell, 174(4):1015–1030.e16.

Schulte, D. and Geerts, D. (2019). Meis transcription factors in development and disease. Development, 146(16).

Selleri, L., Zappavigna, V., and Ferretti, E. (2019). ’building a perfect body’: control of vertebrate organogenesis by pbx-dependent regulatory networks. Genes Dev, 33(5-6):258–275.

Shang, Z., Yang, L., Wang, Z., Tian, Y., Gao, Y., Su, Z., Guo, R., Li, W., Liu, G., Li, X., Yang, Z., Li, Z., and Zhang, Z. (2022). The transcription factor zfp503 promotes the d1 msn identity and represses the d2 msn identity. Front Cell Dev Biol, 10:948331.

Shen, M. W., Arbab, M., Hsu, J. Y., Worstell, D., Culbertson, S. J., Krabbe, O., Cassa, C. A., Liu, D. R., Gifford, D. K., and Sherwood, R. I. (2018). Predictable and precise template-free crispr editing of pathogenic variants. Nature, 563(7733):646–651.

Shen, W. F., Montgomery, J. C., Rozenfeld, S., Moskow, J. J., Lawrence, H. J., Buchberg, A. M., and Largman, C. (1997). Abdb-like hox proteins stabilize dna binding by the meis1 homeodomain proteins. Mol Cell Biol, 17(11):6448–58.

Shohayeb, B., Muzar, Z., and Cooper, H. M. (2021). Conservation of neural progenitor identity and the emergence of neocortical neuronal diversity. Semin Cell Dev Biol, 118:4–13.

Song, X., Chen, H., Shang, Z., Du, H., Li, Z., Wen, Y., Liu, G., Qi, D., You, Y., Yang, Z., Zhang, Z., and Xu, Z. (2021). Homeobox gene six3 is required for the differentiation of d2-type medium spiny neurons. Neurosci Bull, 37(7):985–998.

Squair, J. W., Gautier, M., Kathe, C., Anderson, M. A., James, N. D., Hutson, T. H., Hudelle, R., Qaiser, T., Matson, K. J. E., Barraud, Q., Levine, A. J., La Manno, G., Skinnider, M. A., and Courtine, G. (2021). Confronting false discoveries in single-cell differential expression. Nat Commun, 12(1):5692.

Storm, E. E., Garel, S., Borello, U., Hebert, J. M., Martinez, S., McConnell, S. K., Martin, G. R., and Rubenstein, J. L. R. (2006). Dose-dependent functions of fgf8 in regulating telencephalic patterning centers. Development, 133(9):1831–44.

Stühmer, T., Anderson, S. A., Ekker, M., and Rubenstein, J. L. R. (2002). Ectopic expression of the dlx genes induces glutamic acid decarboxylase and dlx expression. Development, 129(1):245–52.

Su, Z., Wang, Z., Lindtner, S., Yang, L., Shang, Z., Tian, Y., Guo, R., You, Y., Zhou, W., Rubenstein, J. L., Yang, Z., and Zhang, Z. (2022). Dlx1/2-dependent expression of meis2 promotes neuronal fate determination in the mammalian striatum. Development, 149(4).

Su-Feher, L., Rubin, A., Silberberg, S., Catta-Preta, R., Lim, K., Zdilar, I., McGinnis, C., McKinsey, G., Rubino, T., Hawrylycz, M., Thompson, C., Gartner, Z., Puelles, L., Zeng, H., Rubenstein, J., and Nord, A. (2021). Single cell enhancer activity maps neuronal lineages in embryonic mouse basal ganglia. bioRxiv.

Tan, C. L., Plotkin, J. L., Venø, M. T., von Schimmelmann, M., Feinberg, P., Mann, S., Handler, A., Kjems, J., Surmeier, D. J., O’Carroll, D., Greengard, P., and Schaefer, A. (2013). Microrna-128 governs neuronal excitability and motor behavior in mice. Science, 342(6163):1254–8.

Tang, K., Rubenstein, J. L. R., Tsai, S. Y., and Tsai, M.-J. (2012). Coup-tfii controls amygdala patterning by regulating neuropilin expression. Development, 139(9):1630–9.

Teixeira, J. R., Szeto, R. A., Carvalho, V. M. A., Muotri, A. R., and Papes, F. (2021). Transcription factor 4 and its association with psychiatric disorders. Transl Psychiatry, 11(1):19.

Thorvaldsdóttir, H., Robinson, J. T., and Mesirov, J. P. (2013). Integrative genomics viewer (igv): high-performance genomics data visualization and exploration. Brief Bioinform, 14(2):178–92.

Toresson, H., Mata de Urquiza, A., Fagerström, C., Perlmann, T., and Campbell, K. (1999). Retinoids are produced by glia in the lateral ganglionic eminence and regulate striatal neuron differentiation. Development, 126(6):1317–26.

Verheije, R., Kupchik, G. S., Isidor, B., Kroes, H. Y., Lynch, S. A., Hawkes, L., Hempel, M., Gelb, B. D., Ghoumid, J., D’Amours, G., Chandler, K., Dubourg, C., Loddo, S., Tümer, Z., Shaw-Smith, C., Nizon, M., Shevell, M., Van Hoof, E., Anyane-Yeboa, K., Cerbone, G., Clayton-Smith, J., Cogné, B., Corre, P., Corveleyn, A., De Borre, M., Hjortshøj, T. D., Fradin, M., Gewillig, M., Goldmuntz, E., Hens, G., Lemyre, E., Journel, H., Kini, U., Kortüm, F., Le Caignec, C., Novelli, A., Odent, S., Petit, F., Revah-Politi, A., Stong, N., Strom, T. M., van Binsbergen, E., DDD study, Devriendt, K., and Breckpot, J. (2019). Heterozygous loss-of-function variants of meis2 cause a triad of palatal defects, congenital heart defects, and intellectual disability. Eur J Hum Genet, 27(2):278–290.

Visel, A., Minovitsky, S., Dubchak, I., and Pennacchio, L. A. (2007). Vista enhancer browser–a database of tissue-specific human enhancers. Nucleic Acids Res, 35(Database issue):D88–92.

Visel, A., Taher, L., Girgis, H., May, D., Golonzhka, O., Hoch, R. V., McKinsey, G. L., Pattabiraman, K., Silberberg, S. N., Blow, M. J., Hansen, D. V., Nord, A. S., Akiyama, J. A., Holt, A., Hosseini, R., Phouanenavong, S., Plajzer-Frick, I., Shoukry, M., Afzal, V., Kaplan, T., Kriegstein, A. R., Rubin, E. M., Ovcharenko, I., Pennacchio, L. A., and Rubenstein, J. L. R. (2013). A high-resolution enhancer atlas of the developing telencephalon. Cell, 152(4):895–908.

Vogt, D., Hunt, R. F., Mandal, S., Sandberg, M., Silberberg, S. N., Nagasawa, T., Yang, Z., Baraban, S. C., and Rubenstein, J. L. R. (2014). Lhx6 directly regulates arx and cxcr7 to determine cortical interneuron fate and laminar position. Neuron, 82(2):350–64.

Weinreb, C., Rodriguez-Fraticelli, A., Camargo, F. D., and Klein, A. M. (2020). Lineage tracing on transcriptional landscapes links state to fate during differentiation. Science, 367(6479).

Whitington, T., Frith, M. C., Johnson, J., and Bailey, T. L. (2011). Inferring transcription factor complexes from chip-seq data. Nucleic Acids Res, 39(15):e98.

Wonders, C. P. and Anderson, S. A. (2006). The origin and specification of cortical interneurons. Nat Rev Neurosci, 7(9):687–96.

Yang, S., Corbett, S. E., Koga, Y., Wang, Z., Johnson, W. E., Yajima, M., and Campbell, J. D. (2020). Decontamination of ambient rna in single-cell rna-seq with decontx. Genome Biol, 21(1):57.

Yun, K., Garel, S., Fischman, S., and Rubenstein, J. L. R. (2003). Patterning of the lateral ganglionic eminence by the gsh1 and gsh2 homeobox genes regulates striatal and olfactory bulb histogenesis and the growth of axons through the basal ganglia. J Comp Neurol, 461(2):151–65.

Zhang, B., Liu, M., Fong, C.-T., and Iqbal, M. A. (2021). Meis2 (15q14) gene deletions in siblings with mild developmental phenotypes and bifid uvula: documentation of mosaicism in an unaffected parent. Mol Cytogenet, 14(1):58.

Zhang, Z., Gutierrez, D., Li, X., Bidlack, F., Cao, H., Wang, J., Andrade, K., Margolis, H. C., and Amendt, B. A. (2013). The lim homeodomain transcription factor lhx6: a transcriptional repressor that interacts with pituitary homeobox 2 (pitx2) to regulate odontogenesis. J Biol Chem, 288(4):2485–500.

Zhao, L., Liu, Z., Levy, S. F., and Wu, S. (2018). Bartender: a fast and accurate clustering algorithm to count barcode reads. Bioinformatics, 34(5):739–747.

Zhao, Y., Flandin, P., Long, J. E., Cuesta, M. D., Westphal, H., and Rubenstein, J. L. R. (2008). Distinct molecular pathways for development of telencephalic interneuron subtypes revealed through analysis of lhx6 mutants. J Comp Neurol, 510(1):79–99.

Zheng, G. X. Y., Terry, J. M., Belgrader, P., Ryvkin, P., Bent, Z. W., Wilson, R., Ziraldo, S. B., Wheeler, T. D., McDermott, G. P., Zhu, J., Gregory, M. T., Shuga, J., Montesclaros, L., Underwood, J. G., Masquelier, D. A., Nishimura, S. Y., Schnall-Levin, M., Wyatt, P. W., Hindson, C. M., Bharadwaj, R., Wong, A., Ness, K. D., Beppu, L. W., Deeg, H. J., McFarland, C., Loeb, K. R., Valente, W. J., Ericson, N. G., Stevens, E. A., Radich, J. P., Mikkelsen, T. S., Hindson, B. J., and Bielas, J. H. (2017). Massively parallel digital transcriptional profiling of single cells. Nat Commun, 8:14049.

Zolboot, N., Du, J. X., Zampa, F., and Lippi, G. (2021). Micrornas instruct and maintain cell type diversity in the nervous system. Front Mol Neurosci, 14:646072.

Zug, R. (2022). Developmental disorders caused by haploinsufficiency of transcriptional regulators: a perspective based on cell fate determination. Biol Open, 11(1).

